# Internal regulation between constitutively expressed T cell coinhibitory receptors BTLA and CD5 and tolerance in recent thymic emigrants

**DOI:** 10.1101/2024.01.10.574913

**Authors:** Adeolu O. Adegoke, Govindarajan Thangavelu, Ting-Fang Chou, Marcos Petersen, Kiyokazu Kakugawa, Julia F. May, Kevin Joannou, Qingyang Wang, Kristofor K. Ellestad, Louis Boon, Peter A. Bretscher, Hilde Cheroutre, Mitchell Kronenberg, Troy A. Baldwin, Colin C. Anderson

## Abstract

Several coinhibitory receptors are upregulated upon activation, whereas a small number of coinhibitory receptors are expressed constitutively by naive T cells. The relationship between constitutively expressed coinhibitors is unknown. We found an inverse relationship between two constitutively expressed coinhibitors, CD5 and BTLA; BTLA expression was low in the thymus and high in the periphery, corresponding respectively with high and low CD5 expression. Germline or induced deletion of *Btla* in somatic cells demonstrated a causal relationship between BTLA expression and CD5 levels in T cells of central and peripheral lymphoid tissues. The effect of BTLA on CD5 expression on thymic and peripheral CD4 T cells was due to BTLA signaling, rather than signaling by its ligand, the herpes virus mediator (HVEM). Regulation was maintained in mice with a non-signaling HVEM mutant but was lost in *Tnfrsf14*^−/−^ (*Hvem^−/−^*) mice. Increased CD5 levels have been positively associated with increased recognition of self-peptide MHC complexes. Thus, control of CD5 expression by BTLA signals early in T cell ontogeny suggested that BTLA might be important for establishing self-tolerance in newly generated T cells. Consistent with this concept, we found that BTLA, as well as the inducible coinhibitor PD-1, were needed post thymic selection in recent thymic emigrants (RTE) to establish self-tolerance. RTE lacking BTLA caused a multiorgan autoimmune disease whose development required CD4 T cells and MHC class II. Together, our findings identify a negative regulatory pathway allowing constitutively expressed coinhibitory receptors to calibrate their expression in thymic T cell differentiation. Expression of constitutive and induced coinhibitory receptors is needed to establish tolerance in the periphery for RTE.

## Introduction

To generate repertoire diversity, developing T cells in the thymus express receptors generated from random recombination of TCR gene segments [1]. The outcome of this stochastic TCR gene segment rearrangement is the generation of T cells bearing receptors that recognize foreign antigens or self-antigens as agonists, with the latter potentially promoting autoimmune diseases, although varying degrees of beneficial autoreactivity are present in regulatory T cells (Treg) and innate-like T cells, such as natural killer T cells (NKT cells). Positive selection of T cells creates an additional potential danger, as conventional CD4^+^ and CD8αβ^+^ T cells are positively selected based on an ability to recognize a self-peptide–MHC complex. Hence, a critical function of the immune system is to discriminate self from non-self, failure of which will result in detrimental autoimmunity. Thus, most T cells that bind self-peptide–MHC with sufficient affinity need to be made tolerant, and all T cells need to be maintained in a state where they perceive low-affinity interactions with self-peptide–MHC ligand (e.g., the positively selecting peptide) as a ‘tonic’ signal rather than an agonist signal. Although T cell activation and tonic signaling are determined by the interaction of TCR with specific antigenic peptide–MHC complexes, the functional outcome of the T cell response is strongly influenced by costimulatory and coinhibitory signals [2,3]. Such coinhibitory receptors, like programmed cell death protein-1 (PD-1) and cytotoxic T lymphocyte-associated antigen-4 (CTLA-4), are absent from naïve T cells and are upregulated upon activation [4,5]. In contrast, there are a small number of coinhibitory receptors expressed constitutively by naïve T cells, including CD5, B and T lymphocyte attenuator (BTLA), and V-domain immunoglobulin suppressor of T cell activation (VISTA) [6–9].

CD5 is a coinhibitory receptor constitutively expressed on post-selected thymocytes, mature T cells, and a subset of B cells (B1a cells) [6,9–12]. CD5 is expressed as a 67-kDa type I transmembrane glycoprotein (gp), which belongs to the highly conserved superfamily of protein receptors known as the scavenger receptor cysteine rich (SRCR) superfamily [13]. The proposed ligands for CD5 include: CD72 [14], IL-6 [15,16], gp40–80 [17–19], gp150 [20], gp200 [21], IgV_H_ framework region [22], and CD5 itself [23]; however, their physiological relevance and relative importance in interacting with CD5 remains an active area of investigation. CD5 also appears to function without its extracellular domain, questioning whether a ligand for CD5 is relevant. During T cell activation, CD5 is rapidly recruited and co-localizes with the TCR/CD3 complex at the immune synapse, dampening downstream TCR signals [24,25].

BTLA (CD272) is a negative regulator of antigen receptors on B and T cells, dendritic cells, macrophages, and NK cells [7,26–28]. The BTLA cytoplasmic region contains both ITIM and ITSM motifs and a Grb2 recognition motif [7]. Following T cell activation and the interaction of BTLA with its ligand, the herpesvirus entry mediator (HVEM or CD270), the tyrosine residues in ITIM and ITSM of BTLA are phosphorylated and then recruit SH2-containing tyrosine phosphatase 1 (SHP-1) and SHP-2 phosphatases to dampen TCR signaling [27,29]. A T cell can express both BTLA and HVEM, and HVEM interactions with BTLA between cells, in *trans*, or *cis* interactions in which one cell expresses both binding partners, are inhibitory [30]. Consistent with its coinhibitory function, mice lacking BTLA develop systemic autoimmune disease and multiorgan lymphocytic infiltration [31]; they also have increased frequencies of T follicular helper cells in Peyer’s patches and increased IgG and IgA, the latter affecting the homeostasis of gut bacteria [32]. In contrast, *Cd5^−/−^* mice remain relatively healthy, even in late-life [33]. Although constitutively expressed BTLA and CD5 on T cells have unique non-redundant roles, the relationship between these coinhibitors is not well defined. It was recently reported that CD5 expression levels increased in SHP-1 knockout T cells relative to WT T cells [34]. Since BTLA and CD5 exert their inhibitory function through lymphocyte activation-induced recruitment of SHP-1 to their cytoplasmic tail tyrosine residues, these two coinhibitors might have some overlapping or counter-regulatory functions. In addition, CD5 expression is set during T cell development in the thymus and finely adjusted throughout the life of the T cell [35,36], with CD5 surface expression correlating directly with the avidity or signaling intensity of TCR–self-peptide MHC interaction [35,37]. CD5 is expressed most highly on T cells in the thymus [35,38] and on recent thymic emigrants in the periphery [39]. CD5 levels are modified by the autoimmune disease associated H-2^g7^ haplotype [40]. Given the above, CD5 levels might identify stages where the self-reactive potential of the T cell repertoire is greatest. Herein, we explored the relationship between these coinhibitors and found that BTLA broadly controls CD5 expression levels from early in T cell ontogeny and that both BTLA and PD-1 are needed to establish peripheral tolerance at the recent thymic emigrant stage.

## Results

### Negative regulation of CD5 expression by BTLA

To examine the relationship between CD5 and BTLA, we assessed CD5 and BTLA expression in the steady state in both TCRβ^hi^ single positive (SP) T cells in the thymus and splenic T cells from 7-10-week-old C57BL/6 mice (S1A Fig). Consistent with earlier studies [35,38], thymic SP T cells expressed higher levels of CD5 relative to splenic SP T cells (Fig 1A). In contrast, expression of BTLA was lower in the thymic SP T cells relative to splenic SP T cells (Fig 1A). Analysis of the proportion of BTLA^+^ cells among thymic and splenic SP T cells also revealed a reduced frequency of BTLA^+^ SP T cells in the thymus relative to the spleen (Fig 1B). This inverse relationship between BTLA and CD5 was apparent when comparing expression in central vs peripheral lymphoid organs but not within each tissue individually; T cells with the highest BTLA expression in each tissue did not have reduced CD5 expression (S1B Fig). These data suggested that BTLA and CD5 have opposite trajectories of expression as T cells mature.

**Fig 1.**
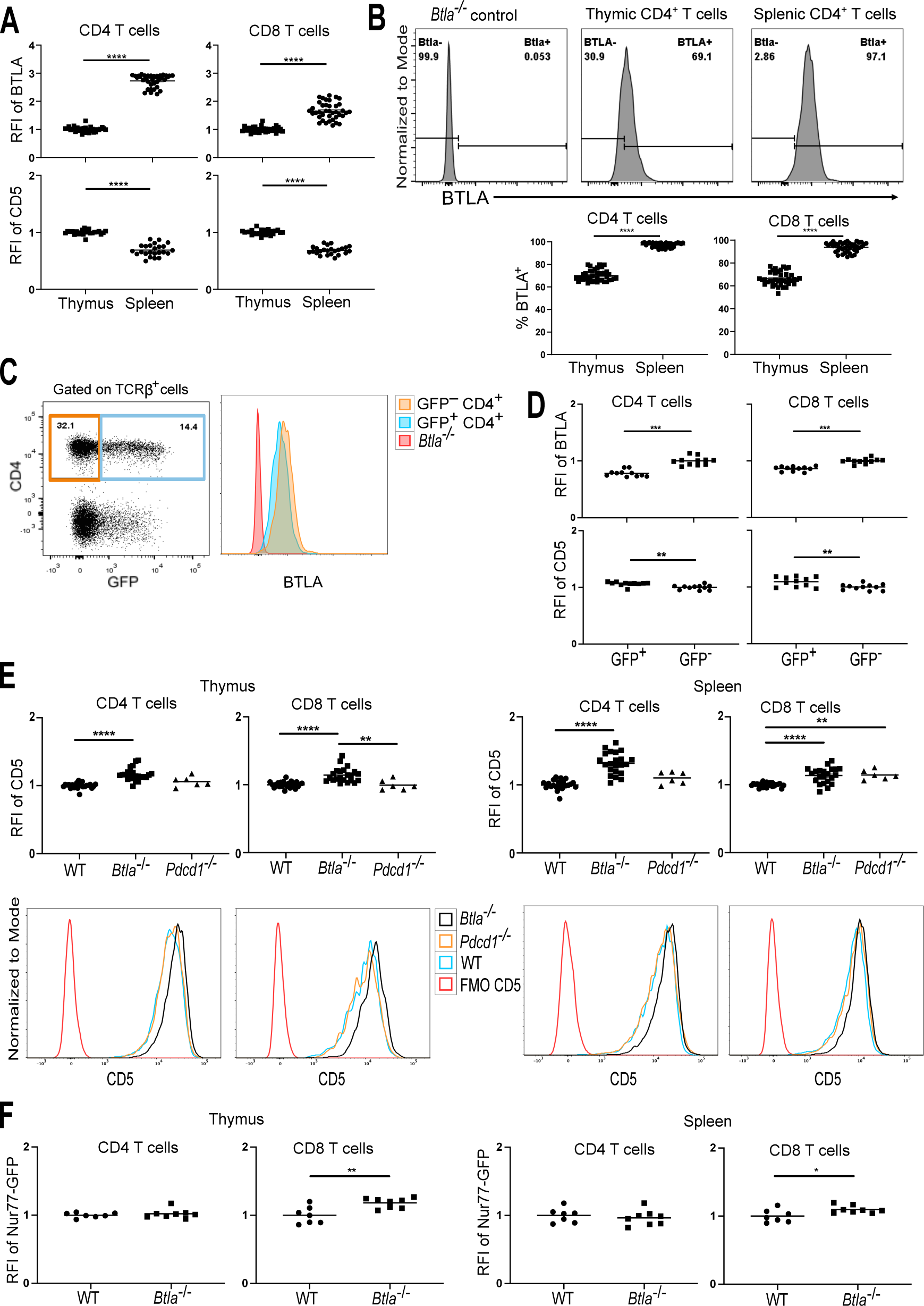
BTLA expression in the thymus and spleen is inversely related to CD5 expression. [**A**] Relative fluorescent intensities (RFIs) of BTLA (upper row) or CD5 (lower row) in CD4 SP T cells (left column) and CD8 SP T cells (right column) in the thymus and spleen (BTLA, n = 39; CD5, n = 24). To calculate the RFI of BTLA or CD5, mean fluorescence intensity (MFI) data were normalized to the average MFI of BTLA or CD5 of the thymic SP T cells in each individual experiment. [**B**] Representative histograms showing the proportion of BTLA^+^ CD4 SP T cells (upper row) in the thymus and spleen. Lower row shows the proportion of BTLA^+^ CD4 SP T cells (lower left) and BTLA^+^ CD8 SP T cells (lower right) in the thymus and spleen (n = 39). [**C**] Representative flow cytometry dot plot (left) and histogram (right) of the indicated markers in splenic TCRβ^+^ cells from 7 - 10-wk old B6.*Rag2p^GFP^* mice. [**D**] RFIs of BTLA (top panels) and CD5 (lower panels) on mature (GFP^−^) or newly generated (GFP^+^) SP T cells in the spleen (n = 11). Data are normalized to the average BTLA MFI or CD5 MFI of the GFP negative splenic SP T cells in each individual experiment. [**E**] CD5 expression on CD4 SP T cells and CD8 SP T cells from thymus (left) or spleen (right) and their corresponding representative histograms (lower rows) from WT (n = 24; B6.*Foxp3^GFP^*, and B6.*Nur77^GFP^*), *Btla^−/−^* (n = 22; B6.*Foxp3^GFP^ Btla^−/−^* and B6.*Nur77^GFP^ Btla^−/−^*), and *Pdcd1*^−/−^ (n = 6; B6.*Foxp3^EGFP^ Pdcd1*^−/−^) mice. [**F**] Nur77-GFP expression. Dots indicate individual mice from six – nine independent experiments (A – B, E) or two independent experiments (C – D, F). \*\**p* < 0.01, \*\*\**p* < 0.001, \*\*\*\**p* < 0.0001.

A small fraction of splenic T cells are recent thymic emigrants (RTE). We hypothesized that these newly generated T cells would also express lower levels of BTLA relative to their established or more mature T cell counterparts. To examine this, we used the *Rag2p^GFP^* mice, where GFP expression is restricted to thymocytes, RTE and newly generated B cells [41,42]. Analysis of BTLA expression comparing the GFP^+^ (RTE) and GFP^−^ (mature) T cells in the spleen revealed that BTLA expression was significantly lower in RTE relative to their mature T cell counterparts (Fig 1C, D); BTLA expression on T cells also increased as GFP expression decreased in both the spleen and thymus (S2 Fig). In contrast, CD5 expression was significantly higher on RTE (Fig 1C, D). Collectively, these data indicated an inverse relationship between CD5 and BTLA expression as T cells mature.

To determine if there was a causal relationship between BTLA expression and CD5 expression, we compared CD5 expression between polyclonal WT and coinhibitor BTLA deficient T cells in the periphery. We analyzed PD-1 in parallel, because we showed earlier that it plays a role in the establishment of tolerance in newly generated T cells [43,44]. The expression of CD5 was significantly higher in *Btla^−/−^* splenic CD4 and CD8 T cells relative to their WT and PD-1 deficient (*Pdcd1^−/−^*) counterparts; the overall increase in CD5 mean fluorescence intensity (MFI) of *Btla^−/−^* T cells was due to a reduction in CD5 low cells (Fig 1E). Congenital absence of the inducible coinhibitor PD-1 resulted in significantly increased CD5 expression in splenic CD8 T cells (Fig 1E). PD-1 deficiency did not alter BTLA expression (S3A Fig). To assess whether the elevated CD5 expression in *Btla^−/−^* splenic T cells originated from the thymus or preferentially increased in the periphery, we compared CD5 expression between the WT, *Pdcd1^−/−^*, and *Btla^−/−^* thymic TCRβ^hi^ SP cells. These cells in the *Btla^−/−^* but not *Pdcd1^−/−^* mice expressed higher CD5 than their WT counterparts (Fig 1E). A possible explanation is that BTLA is expressed on a much higher percentage of thymocytes than PD-1. WT, *Btla^−/−^*, and *Pdcd1^−/−^* T cells expressed a similar level of CD5 in DN and DP populations (S3B, C Fig), indicating that CD5 expression in the *Btla^−/−^* T cells increased only post thymic selection. Although only a small fraction of thymic or splenic T cells expressed the inducible coinhibitor PD-1, BTLA deficiency also increased its expression (S3D Fig).

Treg cells make up a relatively small proportion of the bulk CD4^+^ T cell population, but the preferential expression of higher levels of CD5 in Treg cells [45] may skew the CD5 expression levels in the *Btla^−/−^* CD4^+^ T cells. Therefore, we analyzed Treg numbers [46] and CD5 expression in splenic and thymic TCRβ^+^ CD4^+^ Foxp3^+^ (Treg) or TCRβ^+^ CD4^+^ Foxp3^−^ (non-Treg) cells of WT and *Btla^−/−^* mice (S4 Fig). CD5 expression was significantly higher in both Treg and non-Treg cells of the *Btla^−/−^* mice in the spleen and thymus (S4C, D Fig).

Since CD5 expression is correlated with the signaling strength of TCR:self-antigen (self-pMHC) interactions [35,37] and Nur77 serves as a specific reporter of antigen receptor signaling in murine and human T and B cells [47–53], we hypothesized that Nur77^GFP^ expression would be enhanced in *Btla^−/−^*T cells. We generated *Btla^−/−^ Nur77^GFP^* mice and compared their GFP expression levels in the splenic and thymic T cells to that in BTLA sufficient *Nur77^GFP^* mice. Thymic and splenic *Btla^−/−^*CD8 T cells expressed higher levels of GFP relative to the WT CD8 T cells (Fig 1F). In contrast, GFP expression in the thymic and splenic *Btla^−/−^* CD4 T cells was not significantly different from their WT CD4 T cell counterparts (Fig 1F). Since Nur77^GFP^ expression in B cells also correlates to the B cell receptor affinity for antigen [51], we compared GFP expression levels in B cells from *Btla^−/−^ Nur77^GFP^* mice and WT *Nur77^GFP^* mice and found no significant difference between the two groups (S4E Fig). Together, these data suggest that BTLA expression directly or indirectly determined the level of CD5 expression across the major conventional T cell subsets, while having a more limited effect on Nur77 expression.

### BTLA regulates CD5 expression in adult mice

We determined whether the lack of BTLA from early in T cell ontogeny, as in the case of the germline *Btla* knockout mice, is important for producing heightened T cell CD5 expression. For example, the lack of BTLA during the neonatal period might alter the TCR repertoire when it is first generated, indirectly affecting CD5 levels. We therefore assessed the effect of deleting *Btla* in young adult mice by crossing *B6.Btla^fl/fl^*mice to a tamoxifen-inducible Cre recombinase expressing strain, *B6^Cre/ERT2^*, to generate *B6^Cre/ERT2+/−^ Btla^fl/fl^*mice (S5 Fig). We injected adult 7-week-old *B6^Cre/ERT2+/−^ Btla^fl/fl^* and WT control *B6^Cre/ERT2+/−^* mice intraperitoneally with tamoxifen (Fig 2A). At one-week post-tamoxifen injection only about a third of T cells circulating in the blood had lost BTLA expression. However, the BTLA deficient fraction of CD4 T cells had significantly heightened CD5 expression (Fig 2B). By two weeks post-tamoxifen, *B6^Cre/ERT2+/−^*splenic CD4 and CD8 SP T cells, were 97 ± 2% and 94 ± 4% positive for BTLA, respectively, while *B6^Cre/ERT2+^Btla^fl/fl^*T cells were reduced to 2 ± 1% and 3 ± 2% positive for BTLA (Fig 2C).

**Fig 2.**
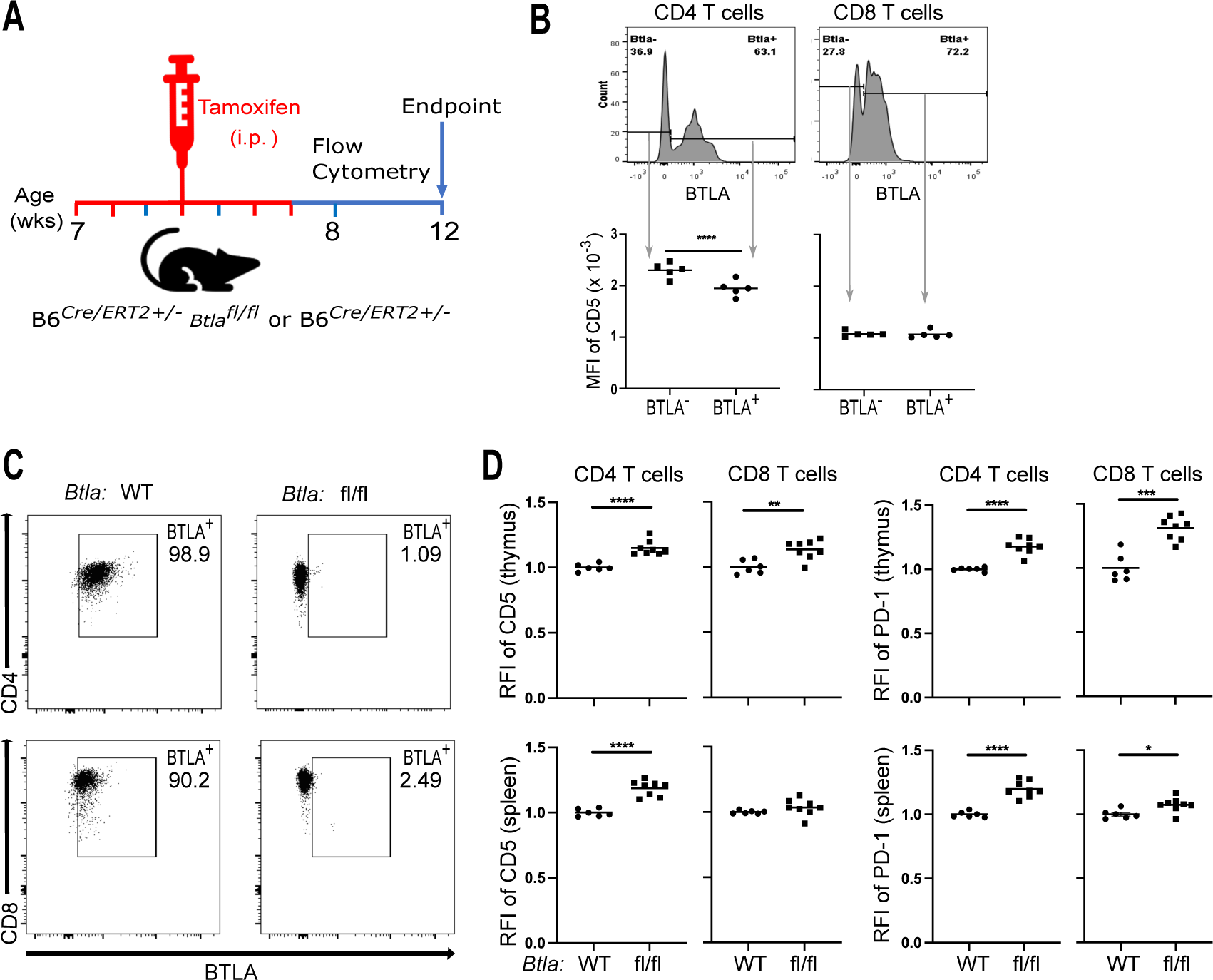
Loss of BTLA in adult mice leads to increased T cell CD5 and PD-1 expression. [**A**] Adult B6*^Cre/ERT2+/−^* or B6*^Cre/ERT2+/−^ Btla^fl/fl^* mice received five doses of tamoxifen on days 0, 1, 3, 5, and 6 (highlighted in red). [**B**] Representative histograms (top) of BTLA expression and the MFI of CD5 (bottom) of BTLA^+^ and BTLA^−^ cells in the CD4 gated (left) and CD8 gated (right) T cells in the peripheral blood at one week post tamoxifen. [**C**] Representative dot plots of BTLA expression in the splenic CD4 SP T cells (top) and CD8 T cells (bottom) in the B6*^Cre/ERT2+/−^* mice (left) and B6*^Cre/ERT2+/−^ Btla^fl/fl^* mice (right) at two weeks post tamoxifen. [**D**] RFI of CD5 and PD-1 in the thymic (top row) CD4 and CD8 or splenic (bottom row) CD4 and CD8 T cells of B6*^Cre/ERT2+/−^* (WT) and B6*^Cre/ERT2+/−^ Btla^fl/fl^* (fl/fl) mice at four weeks post tamoxifen. Dots indicate individual mice from two independent experiments. \*\**p* < 0.01, \*\*\*\**p* < 0.0001.

CD5 expression on thymic and splenic T cells had not yet significantly changed at this time point relative to WT controls (data not shown). By four weeks post-tamoxifen CD5 and PD-1 expression in the thymic and splenic CD4 T cells and thymic CD8 T cells of *B6^Cre/ERT2+/−^ Btla^fl/fl^* mice had increased relative to their *B6^Cre/ERT2+/−^* counterparts (Fig 2D). These data indicated that the impact of BTLA deficiency on CD5 and PD-1 expression is not specific to T cell ontogeny early in life; rather, it can be instigated acutely in adulthood. The rapid increase in CD5 expression on peripheral circulating CD4 T cells, induced by BTLA deletion, suggested that BTLA may be capable of regulating CD5 expression after the early generation of the TCR repertoire and in mature, peripheral T cells. Consistent with the possibility that BTLA regulates CD5 independent of TCR repertoire changes, we did not detect any major differences in the repertoire of sorted thymic CD4 SP T cells of WT and *Btla^−/−^* mice assessed by single cell RNA sequencing (S6 Fig); however, these findings do not rule out potential subtle differences in the repertoire of *Btla^−/−^* mice. Therefore, we tested the effect of BTLA on CD5 expression when the TCR repertoire is fixed. We crossed the *Btla* gene knockout strain to OT-II [54] and OT-I [55] TCR transgenic mice. Complete BTLA deficiency (OT-II.*Btla^−/−^*) led to heightened CD5 expression in OT-II T cells in the periphery and the thymus (Fig 3). OT-II T cells that expressed high levels of the transgenic TCR lacked endogenous TCR expression [56]. The heightened CD5 in BTLA deficient OT-II T cells was evident also in those CD4 SP thymocytes and splenocytes expressing high levels of the transgenic TCR (Vβ5Vα2), indicating that endogenous TCR expression and TCR repertoire changes were not responsible for the altered CD5 expression (Fig 3). In a preliminary analysis of OT-I CD8 T cells, that naturally expressed very high levels of CD5 [57], BTLA deficiency further increased CD5 expression (S7 Fig).

**Fig 3.**
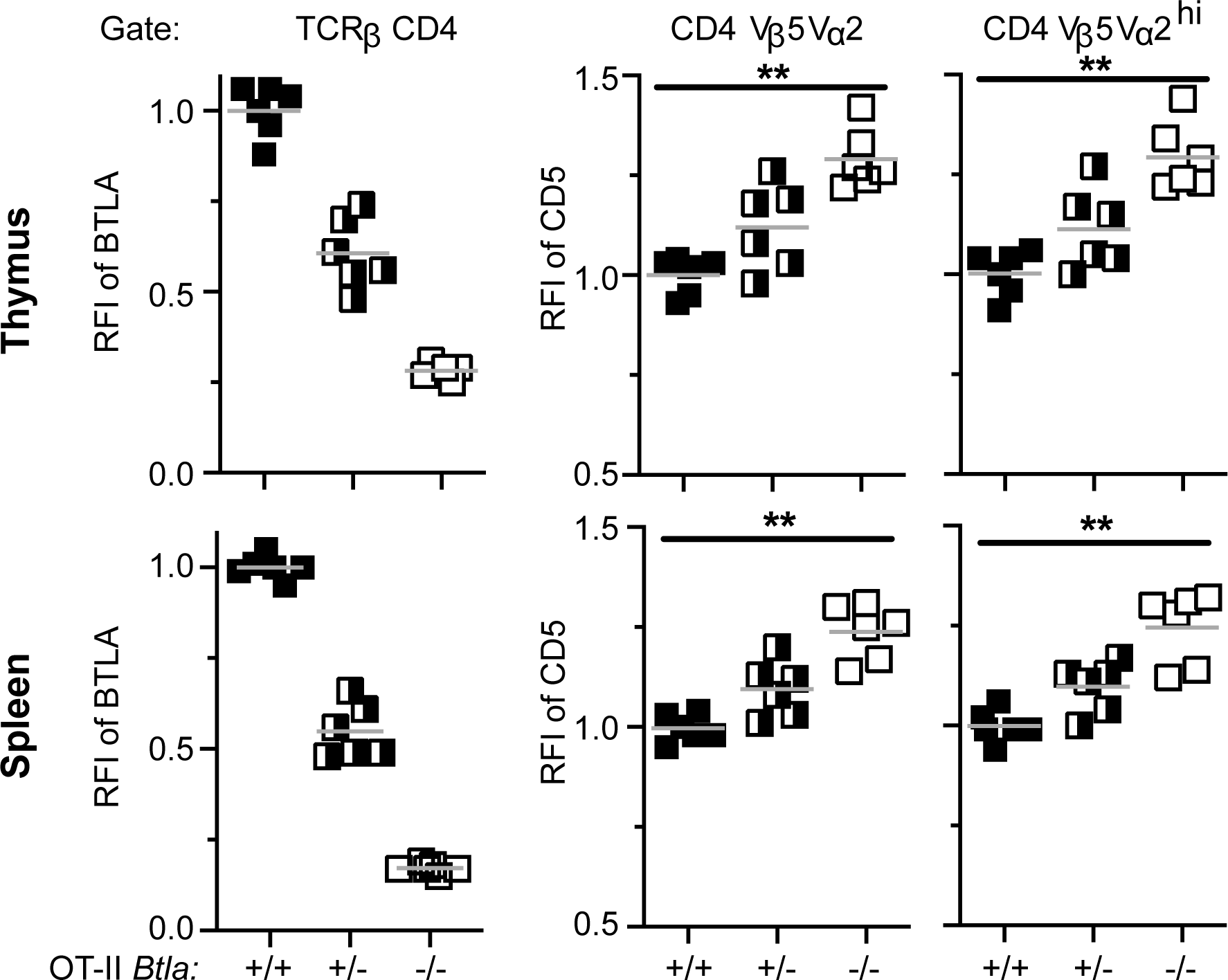
BTLA deficiency increases CD5 levels in TCR transgenic OT-II CD4 T cells in thymus and spleen. RFI of BTLA gated on TCRβ^+^ CD4^+^ SP cells (left) and RFI of CD5 in all CD4^+^ TCR-Vβ5Vα2^+^ cells (middle) and RFI of CD5 in CD4^+^ SP cells expressing high levels TCR-Vβ5Vα2 (right; gate included only the top approximately 50% of TCR-Vβ5Vα2 expressing cells). Analysis of T cells from the thymus (top) and spleen (bottom) of OT-II.*Btla^+/+^* (WT OT-II; n = 6), OT-II.*Btla^+/−^* (n = 6) and OT-II.*Btla^−/−^*mice (n = 6). The grey line is the mean of the RFI. CD5 expression on OT-II.*Btla^+/+^* T cells was significantly lower than on BTLA deficient (*Btla^−/−^*) OT-II T cells, \*\**p* < 0.01.

### HVEM signals do not net set the level of CD5 expression

The only known ligand for BTLA is HVEM, a member of the tumor necrosis factor superfamily (TNFRSF) and therefore is designated as TNFRSF14. HVEM also functions as a signaling receptor, recruiting TRAF2 and activating NK-κβ Rel A, to co-stimulate T cells [58]. HVEM signaling is also involved in survival of memory CD4 [59] and memory CD8 T cells [60,61]. Given the potential for bi-directional signaling, regulation of CD5 and PD-1 expression by BTLA could be due to BTLA signals, HVEM-mediated signals, or both. We therefore examined CD5 and PD-1 expression in germline HVEM knockout mice and mice engineered to express a ‘tail mutant’ HVEM with the intact ectodomain but lacking the normal sequence of the intracytoplasmic domain and therefore incapable of activating NF-κB (*Hvem^tm/tm^*). Using CRISPR-Cas9 technology, we generated an *Hvem* mutant lacking exon 7 with a cytoplasmic tail sequence created by splicing exon 6 to exon 8 (S8A Fig). This sequence lacks the proline-x-glutamate amino acid motif likely required for tumor necrosis factor receptor-associated factor (TRAF) binding and HVEM signaling [62]. We validated the lack of HVEM signaling by the mutant HVEM in vitro in transfected cells using cells expressing HVEM ligands and interacting with cells expressing the tail mutant HVEM with an NF-κB reporter as a readout (S8B Fig). In vivo, the tail mutant HVEM had lower surface expression than either WT mice or *Hvem^WT/tm^* hemizygous mice, (S8C Fig). Despite this, we validated that the reduced amount of surface HVEM was sufficient in vivo in a liver injury model induced by activating invariant natural killer T (iNKT) cells with the potent agonist α-galactosylceramide (αGalCer). Previous studies show that either BTLA or CD160 expressed serve as attenuators of αGalCer-mediated acute hepatic injury [63–65], and this occurs via engagement of HVEM [66]. To test the importance of HVEM signaling function in this hepatic injury model, we co-housed littermate WT control, *Hvem^tm/+^* hemizygous, and *Hvem^tm/tm^*mice that were injected with αGalCer and analyzed 24h later. *Hvem^tm/tm^*mice displayed a similar serum alanine aminotransferase (ALT) level compared to WT or hemizygous mice (S8D Fig), indicating HVEM acts as a ligand for signaling inhibitory receptors in this model.

We found that CD4 SP T cells from *Hvem*^−/−^ mice expressed higher CD5 in the thymus and spleen as well as higher PD-1 in the thymus (Fig 4A). This elevated CD5 and PD-1 expression occurred despite thymic and splenic CD4 and CD8 SP T cells also having heightened BTLA expression in *Hvem*^−/−^ mice, in agreement with a previous report [67]. The heightened BTLA expression, however, could not compensate for the lack of the only known BTLA binding partner. In contrast, despite having lower levels of HVEM than either WT mice or *Hvem^WT/tm^*hemizygous mice, HVEM expression by *Hvem^tm/tm^* mice was sufficient to keep CD5 and PD-1 expression at a level similar to that in WT mice (Fig 4B and S8E Fig). This suggests it is engagement with BTLA that underlies HVEM’s influence on CD5 and PD-1 expression, not signaling through the HVEM cytoplasmic domain.

**Fig 4.**
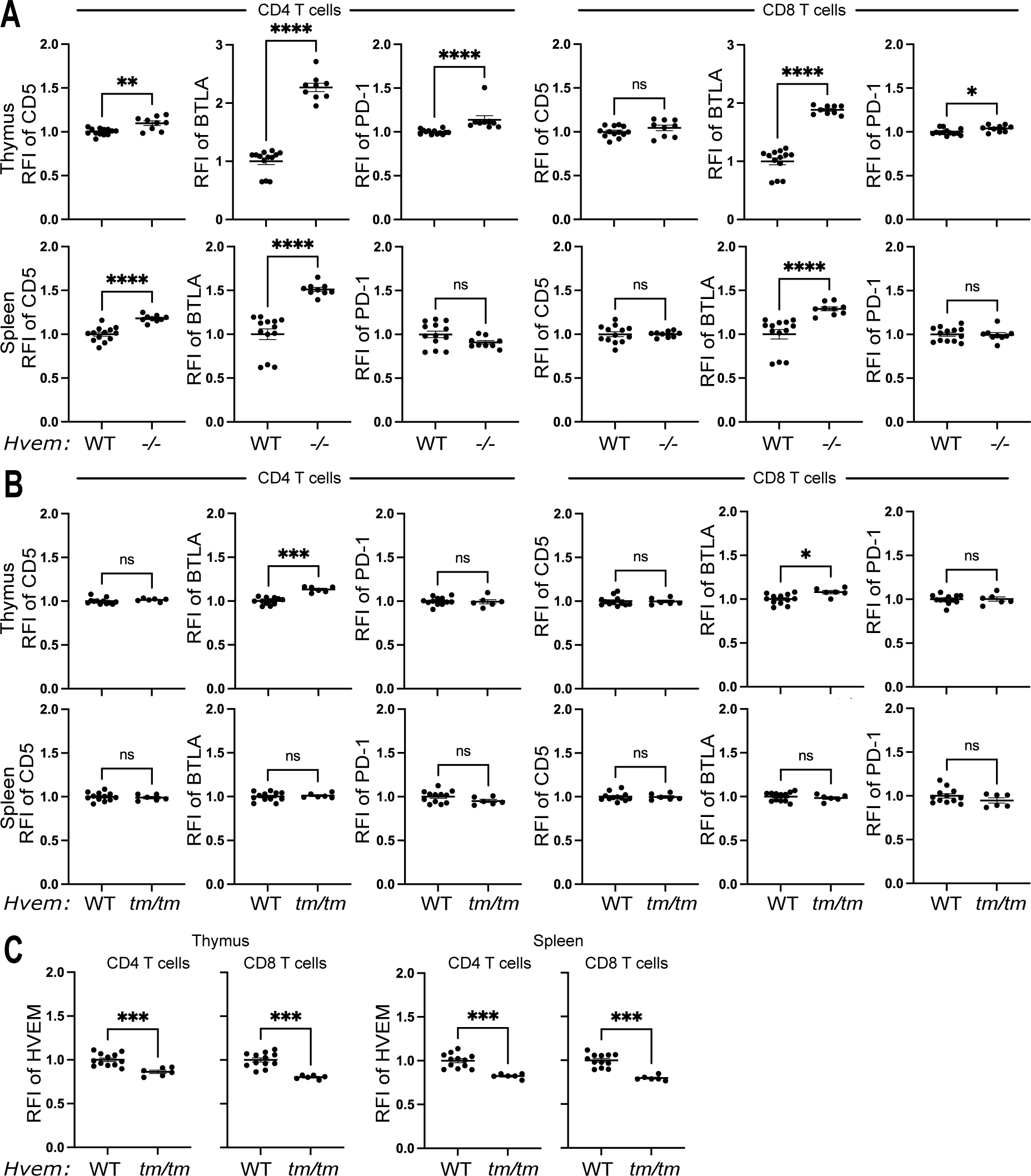
HVEM dependent control of CD5 and PD-1 expression from early in CD4 T cell ontogeny does not require HVEM signaling. Thymocytes and splenocytes from WT and *Hvem^−/−^* [**A**] or *Hvem^tm/tm^*mice [**B, C**], gated on SP CD4 and SP CD8 T cells, were examined by flow cytometry for expression of CD5, BTLA, PD-1 and HVEM. \*\**p* < 0.01, \*\*\**p* < 0.001, \*\*\*\**p* < 0.0001.

### BTLA signaling in newly generated T cells blocks autoimmune disease

Expression of BTLA and its effects on CD5 levels early in T cell ontogeny (Fig 1B, D) may reflect a need for this coinhibitor to ‘tune’ developing T cells to establish tolerance to self-peptide/MHC complexes. Previously, we showed that newly generated T cells depended on PD-1 to broadly establish self-tolerance; transfer of PD-1 deficient fetal liver cells (FLC) to syngeneic *Rag^−/−^* recipients led rapidly to a lethal multiorgan autoimmune disease as newly generated T cells emerged from the thymus [43,44]. To test if BTLA expression is similarly functionally important in newly generated T lymphocytes, we used three approaches that included either stimulating BTLA signaling or removing it. First, we treated recipients of *Pdcd1^−/−^* FLC with an agonistic antibody to BTLA. In this hematopoietic stem cell (HSC) transfer model, T cells typically begin to seed the periphery approximately three weeks post-FLC injection [43]. Shortly after T cell generation recipients of *Pdcd1^−/−^* HSC developed symptoms of disease. The agonistic anti-BTLA antibody significantly delayed disease development (S9A Fig). In the two additional approaches, we adoptively transferred FLC that were either congenitally deficient in BTLA or BTLA was inducibly deleted after transfer (Fig 5A). We transferred FLC from *B6^Cre/ERT2+/−^ Btla^fl/fl^* or *B6^Cre/ERT2+/−^* control mice to adult *Rag^−/−^* mice, followed by injection of tamoxifen to induce *Btla* gene deletion in the transferred cells. FLC was used as a source of HSC in this experiment, allowing for the deletion of BTLA in T cell progenitors pre-thymic selection. Similar to the germline *Btla^−/−^*T cells, peripheral T cells in the recipients of FLC from *B6^Cre/ERT2+/−^ Btla^fl/fl^* mice had increased CD5 expression level through the 8-week experimental period (Fig 5B). T cells were detected in the peripheral blood of FLC recipient mice around four weeks post FLC transfer, which coincided with the onset of disease in recipients of *B6^Cre/ERT2+/−^ Btla^fl/fl^* FLC (Fig 5C). All the recipients of *B6^Cre/ERT2+/−^ Btla^fl/fl^* FLC were diseased before the experimental end-point while their *B6^Cre/ERT2+/−^*counterparts remained free of disease (Fig 5C). Lymphocytes populating the periphery of recipients of *B6^Cre/ERT2+/−^ Btla^fl/fl^* FLC were BTLA negative while lymphocytes populating *B6^Cre/ERT2+/−^* FLC recipients expressed BTLA (Fig 5D). Although BTLA expression has been seen on non-hematopoietic cells in some settings [68,69], these data indicated that loss of BTLA in fetal liver derived cells alone was sufficient for the generation of multi-organ autoimmune disease in the recipients.

**Fig 5.**
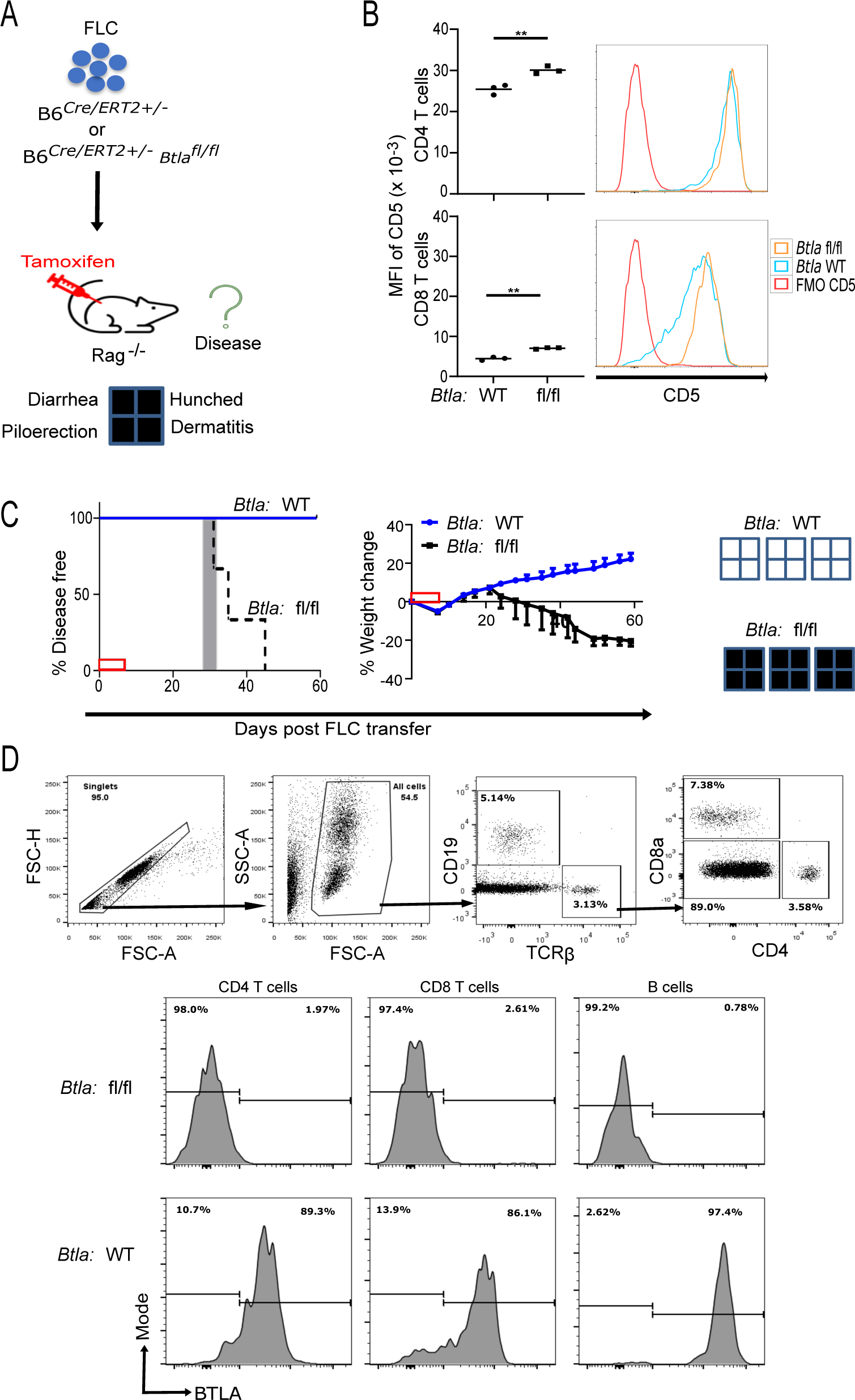
Loss of BTLA early in T cell ontogeny generates autoimmune disease. [**A**] We adoptively transferred 20 x 10^6^ FLC pooled from 8-10 embryonic day 14 - 16 B6*^Cre/ERT2+/−^* (WT) or B6*^Cre/ERT2+/−^ Btla^fl/fl^* (fl/fl) fetuses to 7-wk old *Rag^−/−^*mice on day 0 (n = 3 recipients per group), followed by tamoxifen injection on days 0, 1, 3, 5, and 6. Recipient mice were monitored for signs of disease for eight weeks post FLC transfer. [**B**] MFI of CD5 in CD4 T cells (top) and CD8 T cells (bottom) with respective representative histograms of peripheral T cells in the recipients of FLC from B6*^Cre/ERT2+/−^* and B6*^Cre/ERT2+/−^ Btla^fl/fl^* at eight weeks post tamoxifen. Dots indicate data from individual mice; \*\**p* <0.01. [**C**] Left panel: disease incidence in recipients of B6*^Cre/ERT2+/−^* (blue line) B6*^Cre/ERT2+/−^ Btla^fl/fl^*(black dashed line) FLC. Survival curves were significantly different, *p* = 0.02. The grey rectangle indicates the range, in days, at which the first T cells were detected in the peripheral blood after FLC transfer. Right panel: weight changes in recipients of B6*^Cre/ERT2+/−^ Btla^fl/fl^* FLC or B6*^Cre/ERT2+/−^* FLC. The red box on the X-axes indicates the tamoxifen treatment period. The presence (filled) or absence (empty) of disease signs is depicted on the far-right panel. [**D**] Flow cytometry gating (top). A representative histogram of BTLA expression in the T and B cells populating the periphery of B6*^Cre/ERT2+/−^ Btla^fl/fl^*FLC recipient mice (middle) or B6*^Cre/ERT2+/−^ Btla^fl/fl^* FLC recipient mice (bottom) at four weeks post FLC transfer is shown.

To examine in more detail the autoimmune disease generating effects of BTLA deficiency in newly generated vs. established *Btla^−/−^*T cells, we compared transfers of FLC, thymocytes, whole splenocytes or sorted splenic T cells to syngeneic *Rag^−/−^* recipients. Most of the recipients of *Btla^−/−^* thymocytes or FLC demonstrated severe morbidity, while recipients of the WT control FLC, FLC deficient in Fas (*lpr* FLC), and recipients of *Btla^−/−^* whole splenocytes or purified splenic T cells remained free of disease (Fig 6A). Histological analysis of tissue sections obtained from sick mice 60-65 days post cell transfer revealed lymphocytic infiltrations in major organs, including the liver, kidney, and pancreas of *Btla^−/−^*FLC recipients (S9B Fig). Lymphocytic infiltration was present in the liver of *Btla^−/−^* FLC but not WT FLC recipients and included CD4 and CD8 T cells (Fig 6B). The liver was the most frequently affected internal organ examined in *Rag^−/−^*recipients of *Btla^−/−^* FLC, consistent with the late life spontaneous hepatitis that has been observed in unmanipulated *Btla^−/−^*mice [31]. Newly generated *Btla^−/−^* T cells demonstrated substantially increased proliferation compared to WT T cells (Fig 6C). Collectively, these data demonstrated a requirement for BTLA in newly generated T cells to establish tolerance and prevent lymphopenia-potentiated autoimmune disease, while BTLA was not required for maintaining tolerance of established T cells under these conditions.

**Fig 6.**
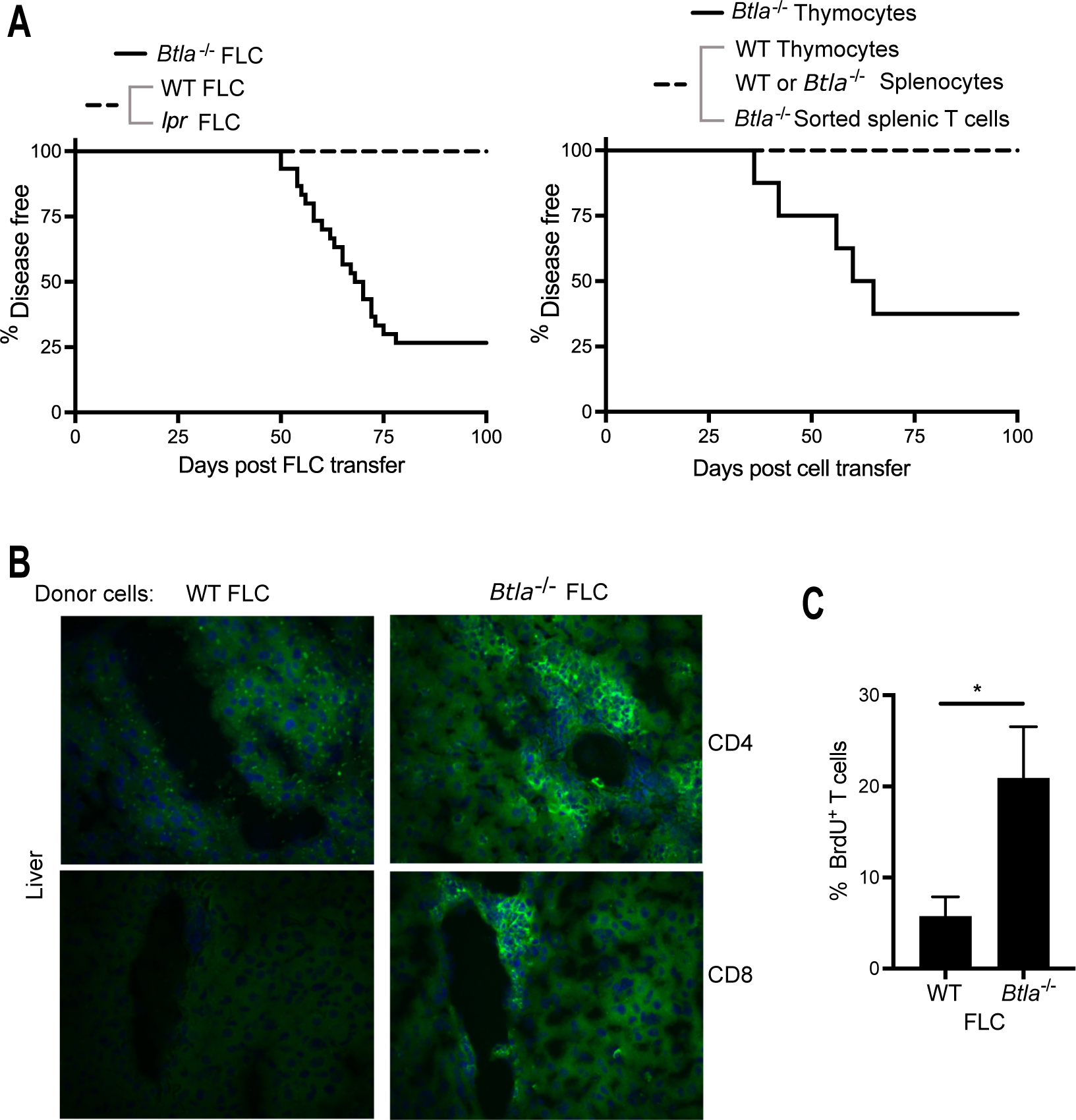
BTLA is needed for establishment of tolerance in newly generated T cells. [**A**] *Left:* Adult *Rag^−/−^*mice were given fetal liver cells from *Btla^−/−^* (n = 30) or WT (n = 10) or *lpr* fetuses (n = 3) and disease was monitored. *Right:* Adult *Rag^−/−^* mice were given one of the following cell populations (all populations contained 2-2.5 x 10^6^ single positive T cells), *Btla^−/−^* (n = 8) or WT thymocytes (n=4), or WT or *Btla^−/−^* splenocytes (n = 4), or FACS sorted *Btla^−/−^* splenic TCR^+^ CD24^low^ T cells (n = 3) and were monitored for disease incidence (diseased mice exhibited two or more of the following symptoms: loss of weight, hunched posture, piloerection, dermatitis and diarrhea.). Disease was only observed in *Btla^−/−^* FLC or thymocyte groups, with all other groups surviving to day 100 disease-free. [**B**] Representative immunofluorescence (n = 5-6/group; original magnification × 400) of liver from individual recipients of WT or *Btla^−/−^* FLC. Blue: staining with the nuclear marker 4’,6’-diamidino-2-phenylindole (DAPI); green: CD4 or CD8 staining. [**C**] The mean percentage of BrdU^+^ T cells in the spleen of *Rag^−/−^* recipients (n = 4/group) of *Btla^−/−^*FLC was significantly higher (\**p* < 0.05) than that of WT FLC recipients examined 65 days post-FLC transfer.

### CD4^+^ T cells and MHC II are required for autoimmune disease

Having shown that loss of BTLA in newly generated T cells leads to autoaggressive T cells, we asked what T cell subset(s) is required for disease generation. We adoptively transferred sorted CD4 or CD8 SP thymocytes from *Btla**^−/−^*** or WT mice into *Rag**^−/−^***mice (Fig 7A and S10A Fig). We monitored the recipient mice for several weeks or until losing ≥ 20% of baseline body weight, whichever came first. Recipients of *Btla**^−/−^***CD4 SP T cells started losing weight as early as 21 days post cell transfer, which continued for up to 53 days when they had lost ≥ 20% of their baseline body weight (Fig 7A). All of the *Btla**^−/−^*** CD4 SP T cell recipient mice had a hunched appearance, piloerection, diarrhea and some had dermatitis. Disease occurred in male and female *Rag**^−/−^*** recipients of *Btla**^−/−^*** CD4 SP T cells. In contrast, none of the recipients of *Btla**^−/−^*** CD8 SP T cells showed signs of morbidity, and their body weight increased throughout the experiment. Although recipients of WT SP CD4 or CD8 T cells had an initial decline in body weight, they quickly recovered and showed an improvement in body weight and no signs of disease (Fig 7A). To examine what MHC molecule is required for this autoimmune disease, we adoptively transferred whole *Btla**^−/−^*** SP thymocytes to *K^b^D^b**−/−**^ Rag**^−/−^*** or *CiiTA**^−/−^** Rag**^−/−^*** mice that lacked both Rag and MHC class I genes or lacked both Rag and MHC class II protein expression [70], respectively. All *CiiTA**^−/−^**Rag**^−/−^*** mice were free of disease, while all *K^b^D^b**−/−**^ Rag**^−/−^***recipients had signs of morbidity within 3 weeks post T cell transfer, regardless of sex (Fig 7B). Similarly, sorted CD4 *Btla**^−/−^***SP thymocytes transferred to *K^b^D^b**−/−**^ Rag**^−/−^*** caused disease, while sorted CD8 *Btla**^−/−^*** SP thymocytes transferred to *CiiTA**^−/−^** Rag**^−/−^*** mice did not (S10B, C Fig). Thus, MHC class II and MHC class II-restricted CD4 T cells were required to induce disease.

**Fig 7.**
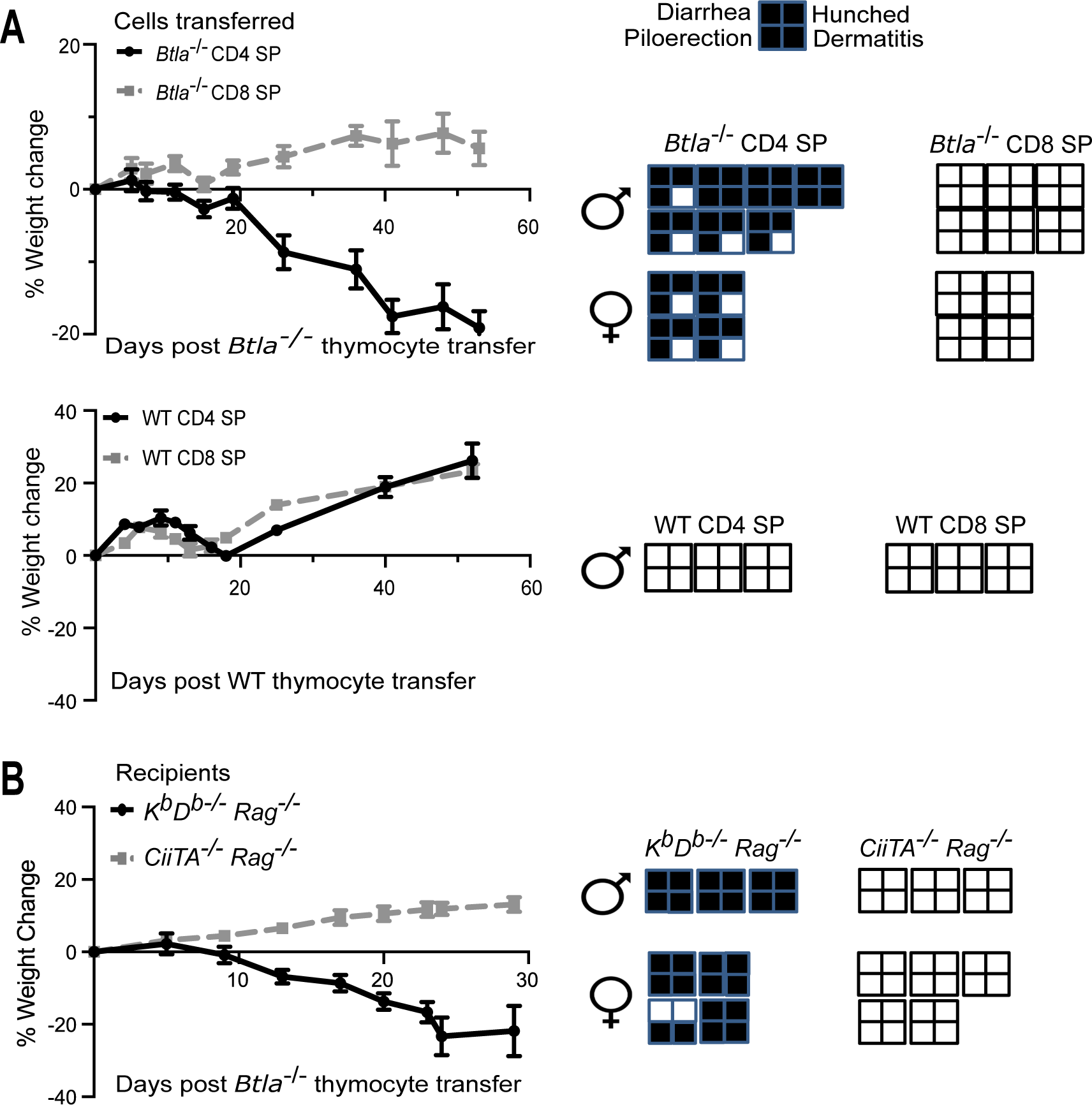
Autoimmune disease in Btla^−/−^ thymocyte recipients requires CD4^+^ T cells and MHC II. [**A**] We adoptively transferred 3 x 10^6^ MACS-sorted CD4 or CD8 SP thymocytes pooled from seven 8 – 10-wk old B6.*Foxp3^EGFP^* x *Btla^−/−^*(left column) or B6.Foxp3^EGFP^ (right column) mice i.v. to 8 – 10-wk old *Rag^−/−^* mice (*Btla^−/−^* thymocyte recipients, n = 10-11 mice/group; WT thymocyte recipients, n=3 mice/group) and monitored for several weeks or after losing ≥ 20% of baseline body weight, whichever came first. Body weight change of the *Btla^−/−^* SP thymocyte recipients (data was from three independent experiments) or WT SP thymocyte recipients is shown, and the presence (shaded) or absence (unshaded) of disease signs is depicted to the right of the graphs. [**B**] Thymocytes containing 3 x 10^6^ SP cells (i.e., non-sorted) pooled from seven 8 – 10-wk old B6.*Foxp3^EGFP^* x *Btla^−/−^* mice were injected via tail vein to 8 – 10-wk old *K^b^D^b–/–^ Rag^−/−^* mice (n = 7) or *CiiTA^−/−^ Rag^−/−^* (n = 8) mice, which were then monitored for several weeks or until after losing ≥ 20% of baseline body weight, whichever came first. Body weight change of recipient mice is shown. Data are from two independent experiments. The presence (shaded) or absence (unshaded) of disease signs is depicted to the right of the graph. Cell donors and recipients were of the sex indicated.

### BTLA and PD-1 signaling are needed in newly generated T cells to block autoimmune disease

The current studies identified that, like PD-1, BTLA was important for the establishment of tolerance in newly generated T cells; however, they did not address whether these coinhibitory receptors were needed during thymic selection or post thymic selection, when the newly generated T cells seed the periphery. To examine if autoimmune disease occurs when BTLA or PD-1 is deleted post thymic selection, thymocytes from adult (7 – 9-week-old) *B6^Cre/ERT2+/−^ Btla^fl/fl^* or *B6^Cre/ERT2+/−^ Pdcd1^fl/fl^* mice were adoptively transferred via the tail vein to adult *Rag**^−/−^*** mice, followed by tamoxifen injection to induce gene deletion in the transferred thymocytes. The control group received *B6^CreERT2+/−^* thymocytes and tamoxifen injection (Fig 8A). There was a near complete deletion of BTLA and PD-1 by day 7 post the last dose of tamoxifen in the T cells of *B6^Cre/ERT2+/−^ Btla^fl/fl^*and *B6^Cre/ERT2+/−^ Pdcd1^fl/fl^* thymocyte recipients, respectively, while expression of these coinhibitors was maintained in the *B6^CreERT2+/−^* thymocyte recipients (Fig 8B). All of the *B6^Cre/ERT2+/−^ Btla^fl/fl^* and *B6^Cre/ERT2+/−^ Pdcd1^fl/f^* thymocyte recipients, but not their *B6^CreERT2+/−^*counterparts, developed inflammatory disease. They began to lose body weight at days 7 - 14 post thymocyte transfer and showed additional signs of disease at days 12 - 17 (Fig 8C). Histological examination showed CD4 SP T cell infiltration in the kidney and liver of *B6^Cre/ERT2+/−^ Btla^fl/fl^* thymocyte recipients and the kidney, liver, and lung of *B6^Cre/ERT2+/−^ PD-1^fl/fl^* thymocyte recipients (S11 Fig). Since BTLA and PD-1 were present during thymic selection and deleted only post cell transfer, these coinhibitors were needed at the RTE stage to establish tolerance.

**Fig 8.**
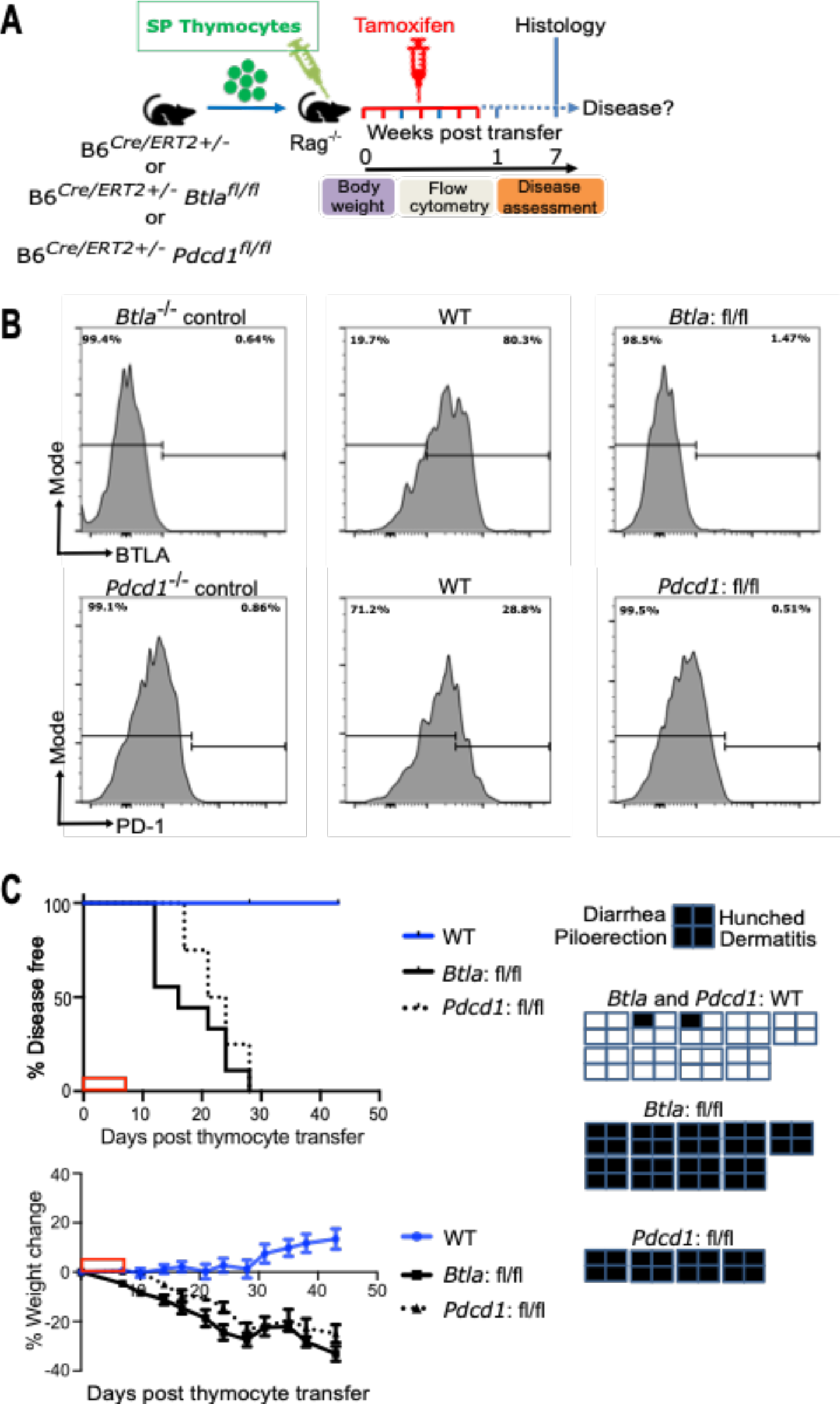
BTLA and PD-1 are needed post thymic selection in newly generated T cells to block autoimmune disease. [**A**] We adoptively transferred thymocytes containing 5 x 10^6^ SP (non-pooled) from 7 – 12-week-old B6*^Cre/ERT2+/−^* (WT) or B6*^Cre/ERT2+/−^ Btla^fl/fl^* (fl/fl) mice or B6^C*re/ERT2+/−*^ *Pdcd1^fl/fl^*(fl/fl) mice to 7 - 12-week-old *Rag^−/−^* mice on day 0, followed by tamoxifen injection on days 0, 1, 3, 5, and 6. Mice were then monitored for signs of disease for seven weeks. [**B**] Flow cytometry gated on TCRβ^+^ cells. Representative histograms of BTLA (top row) or PD-1 (lower row) expression in the T cells of germline gene knockout control mice and B6*^Cre/ERT2+/−^*or B6*^Cre/ERT2+/−^ Btla^fl/fl^*or B6*^Cre/ERT2+/−^ Pdcd1^fl/fl^* thymocyte recipients at two weeks post thymocyte transfer (i.e., seven days post tamoxifen) is shown. [**C**] Top Left panel: disease incidence in recipients of B6*^Cre/ERT2+/−^* (n = 9) or B6*^Cre/ERT2+/−^ Btla^fl/fl^* (n = 9) or B6*^Cre/ERT2+/−^ Pdcd1^fl/fl^* (n = 4) thymocytes. Disease free survival curve comparison demonstrated a significant difference between both fl/fl groups and the WT, with *p* < 0.0001. Data are combined from three independent experiments (two for *Btla*: fl/fl and one for PD-1: fl/fl; WT were included in all three experiments). Bottom left panel: weight changes in the indicated of B6*^Cre/ERT2+/−^ Btla^fl/fl^* or B6*^Cre/ERT2+/−^ Pdcd1^fl/fl^* or B6*^Cre/ERT2+/−^* thymocyte recipients. The presence (shaded) or absence (unshaded) of disease signs is depicted on the far-right panels. The red box on the X-axis indicates the tamoxifen treatment period.

## Discussion

In this study, we aimed to understand the relationship between the constitutively expressed T cell coinhibitory molecules BTLA [71,72] and CD5, and an induced coinhibitor, PD-1, in the steady state and under conditions that promote multi-organ autoimmune disease. Our data on steady state CD5 expression in the thymus and spleen were consistent with findings in the literature [35,38] and when compared with BTLA expression, revealed an inverse relationship. Our data showed that peripheral T cells expressed higher levels of BTLA than thymocytes. This is a result of a maturation process that began in the thymus and continued in the periphery, with RTE in the spleen expressing lower levels of BTLA relative to the mature splenic T cells. In contrast, splenic RTE expressed higher levels of CD5 relative to their mature T cell counterparts, as seen previously [39]. High CD5 on RTE may reflect increased sensitivity to self-peptide-MHC complexes, consistent with the greater self-reactivity and autoimmune potential of RTE that we have observed in the context of PD-1 deficiency [44].

Analysis of CD5 expression in the *Btla^−/−^* mice or upon induced BTLA deletion showed that the overall mean CD5 expression was enhanced in BTLA deficient CD4 and CD8 T cells, indicating that BTLA negatively regulates CD5 expression. Heightened CD5 expression was a result of a reduction in CD5 low cells. This suggests that BTLA was either needed for survival of CD5 low cells or it enhanced CD5 expression in the CD5 low cells. While a role for BTLA in T cell survival during chronic stimulation has been shown [73], we found increased expression of CD5 on BTLA deficient OT-II CD4 T cells, a T cell that normally expresses relatively low CD5 levels [37]. This indicated BTLA may be able to negatively regulate CD5 expression independently from any effects it might have on the generation of the TCR repertoire early in life. CD5 expression on differentiating thymocytes reflects the TCR affinity for self-peptide MHC complexes. Regulation of CD5 expression in thymocytes appeared to be specific to BTLA, as a deficiency in another coinhibitor (PD-1) had no effect on CD5 expression. Complementing the CD5 expression data, the *Btla^−/−^* thymic and splenic CD8 T cells displayed enhanced Nur77^GFP^ expression. Although our data showed a disparity between CD5 and Nur77^GFP^ expression levels in the *Btla^−/−^*thymic and splenic CD4 T cells, Zinzow-Kramer *et al*., had previously reported that a broad range exists in the expression levels of CD5, Ly6C, and Nur77^GFP^ in vivo for naïve T cells even within the same TCR niche [74]. This suggests that CD5 and Nur77 can be regulated differently from each other in response to TCR signals. Heightened PD-1 expression in CD4 and CD8 SP thymocytes in mice deficient for either BTLA or HVEM further supported the concept that loss of BTLA signaling leads to enhanced self-recognition early in T cell ontogeny. Strikingly, even the decreased levels of HVEM expression, as occurred in *Hvem^tm/tm^*mice, were sufficient to negatively regulate both BTLA and PD-1. Given that HVEM interacts with several different ligands, it could not be anticipated whether HVEM deficiency would mimic BTLA deficiency in the regulation of CD5 and PD-1 expression. HVEM deficiency mimicked BTLA deficiency closely, however, for control of CD5 expression in CD4 T cells but not in CD8 T cells. This suggested the other ligands for HVEM, including LIGHT and CD160, may have differential effects on CD5. Together these findings suggest that expression of some co-inhibitors are calibrated by BTLA signals triggered through a highly sensitive engagement by HVEM, acting not as a signaling receptor but purely as a ligand or binding partner for BTLA.

Mice deficient in BTLA develop autoimmune disease only later in life [31], suggesting that either other coinhibitors compensate for the loss of BTLA in early life, or early events take time for their consequences to become apparent. Increased CD5 expression in the *Btla^−/−^*mice suggested that CD5 may serve as a compensatory coinhibitory receptor. Despite this, increased CD5 on newly generated T cells was not sufficient for preventing autoimmune disease following transfer to immune deficient mice. The conditions in *Rag^−/−^* recipients, however, in terms of the extent or rapidity of lymphopenia induced proliferation, changes in the microbiome, and others, might not fully represent the conditions in perinatal immune competent mice. Compensatory coinhibitor expression may limit T cell autoreactivity in other contexts. In cancer therapy it can limit the efficacy of treatment [75]. BTLA blockade has been shown to limit tumour growth and improve survival in a murine model [76] and it is a target in clinical trials [77]. The effect of blocking both CD5 and BTLA have yet to be determined. Supporting our findings in the *Pdcd1*^−/−^ mice, where CD5 expression was elevated in the splenic CD8^+^ T cells, a recent report showed that blocking CD5 together with PD-1 substantially enhanced survival and tumour cell killing [78].

We examined the role of BTLA in newly generated T cells and found that BTLA, like PD-1 [43,44], was important for establishing tolerance in newly generated, polyclonal T cells. In contrast, BTLA signaling appears to be dispensable for establishing tolerance in newly generated TCR transgenic CD8 T cells that recognize a tissue-specific antigen expressed in the thymus and pancreas [79]. This might reflect a more critical role for BTLA in CD4 T cells. Consistent with this idea, we found that sorted *Btla**^−/−^*** CD4 but not CD8 T cells could generate multi-organ autoimmune disease, and disease development did not occur in MHC class II deficient recipients. BTLA was also previously reported to regulate CD4 T cell alloreactivity and proliferative responses to MHC class II antigens [80]. CD4 T cells and/or MHC class II are also required for disease development caused by newly generated *Pdcd1**^−/−^***T cells [43,44].

To understand at what point during T cell development BTLA and PD-1 may be needed to establish tolerance, we deleted each of these coinhibitors individually in RTE that had the capacity to express them during thymic development. We found that loss of BTLA or PD-1 post thymic selection in RTE resulted in autoimmune disease in lymphopenic host mice, associated with loss of weight, dermatitis, diarrhea, and T cell infiltration of organs. Thus, BTLA and PD-1 were needed post thymic selection to establish peripheral tolerance in newly generated T cells, at least in *Rag^−/−^* mice. Our current study does not exclude a role for these coinhibitors during central tolerance. A role for BTLA in central tolerance has not yet been examined. We previously found that PD-1 was not needed for central tolerance in CD4 or CD8 T cells specific to a ubiquitous self-antigen [43], however, PD-1 was needed for non-deletional central tolerance in CD8 T cells that recognize a tissue-restricted antigen [79]. The current data suggest that any role the coinhibitors might have in inducing central tolerance is not by itself sufficient to broadly establish a durable self-tolerance. The need for these coinhibitors in RTE suggests that BTLA and PD-1 will be critical to establishing tolerance during the neonatal period when all peripheral T cells are RTE [81,82]. Consistent with this idea, a substantial proportion of neonatal CD4 T cells, those undergoing natural lymphopenia-induced-proliferation, were found to express PD-1 [44]. Our current studies are further addressing this important question.

Interestingly, adult mouse splenic T cells from mice with germline BTLA or PD-1 [43] deficiency did not cause overt autoimmune diseases in the recipients. From *Pdcd1**^−/−^*** splenocytes, only the sorted RTE subpopulation could generate overt disease; sorted established T cells did not cause disease [44]. A key question for future studies is whether the seeming lack of need for these coinhibitors in established T cells indicates that the coinhibitors are not required to maintain a tolerant state to tonic self-peptide/MHC ligands. An alternative possibility is that these coinhibitors are needed to maintain tolerance when T cells have developed in their presence. Although our study establishes the importance of BTLA and PD-1 signaling in RTE to block autoimmune disease development in lymphopenic recipients, the germline BTLA or PD-1 knockout mice seem to be protected from overt autoimmune disease during the neonatal period when there is physiological lymphopenia. This is likely to result from two factors: (1) their T cell repertoire is first generated early in life, a period naturally deficient in lymph node stroma [83], and (2) early disease-causing events normally take time for their consequences to become apparent. Consistent with the first idea, we previously showed that neonatal *Rag**^−/−^***recipients of *Pdcd1**^−/−^*** FLC were relatively resistant to disease, as were adult recipients that lacked or had reduced lymph nodes [43]. Supporting the second idea, T cells generated early in life in the NOD mice play an important role in the initiation of insulitis long before the onset of overt diabetes that manifests much later in life [84,85].

Taken together, our data identify a regulatory axis between BTLA and CD5 and a critical need for both constitutive and inducible coinhibitors (BTLA and PD-1) when T cells initially seed the periphery.

## Materials and methods

### Animals

Mice used in this study included male and female B6.129S7-*Rag1*^tm1Mom^/J (Rag^−/−^), B6.Cg-*Foxp3*^tm2(EGFP)Tch^/J (*Foxp3^EGFP^*) (34,35), B6.129-Gt(ROSA)26Sor^tm1(cre/ERT2)Tyj^/J (B6*^Cre/ERT2+^*), C57BL/6J (H-2^b^; B6), B6.129S2-Ciita^tm1Ccum^/J (*CiiTA^−/−^*), and B6.MRL-*Tnfrsf6^lpr^*/J (*lpr*) mice were originally purchased from The Jackson Laboratory (Bar Harbor, ME, USA). The C57BL/6 H-2K^btm1^-H-2D^btm1^N12 (*K^b^D^b**−/−**^*) were originally obtained from the National Institute of Allergy and Infectious Diseases (NIAID) Exchange Program (NIH: 004215) [86]. We generated the *CiiTA**^−/−^** Rag**^−/−^*** mice and *K^b^D^b**−/−**^ Rag**^−/−^*** mice by crossing the *Rag^−/−^* mice with *CiiTA**^−/−^*** mice and *K^b^D^b**−/−**^* mice, respectively. B6.*Rag2p^GFP^* mice [41,42] were kindly provided by Dr. Pamela Fink (University of Washington, Seattle, WA), and we previously generated B6.*Rag2p^GFP^ Pdcd1^−/−^* mice [44]. C57BL/6-*Btla^−/−^*(abbreviated as *Btla^−/−^*) were originally provided by Dr. Kenneth Murphy (Washington University, St. Louis, MO), and C57BL/6-*Pdcd1^−/−^*(abbreviated as *Pdcd1^−/−^*; backcrossed 11 generations to C57BL/6; originally generated by Prof. T. Honjo and colleagues [87]) mice were bred at the University of Alberta. We crossed OT-II mice (kindly provided by Dr Sue Tsai, University of Alberta) and OT-I mice (kindly provided by Dr. M. McGargill, St. Jude Children’s Research Hospital, Memphis, TN) to *Btla^−/−^* mice, and crossed *Foxp3^EGFP^* mice to *Btla^−/−^* and *Pdcd1^−/−^* mice [46]. The B6.*Pdcd1^fl/fl^* mice were purchased from Taconic Biosciences (Rensselaer, NY, USA). B6.*Btla^fl/fl^* mice [32,88] generated by us (M.K.) and were kindly provided by John R Šedý and Carl Ware (Sanford Burnham Prebys Medical Discovery Institute, La Jolla, United States). B6.*Nur77^GFP^* mice [53] were provided by Dr Kristin Hogquist (University of Minnesota, Minneapolis, MN) and we generated *Btla^−/−^ Nur77^GFP^* mice. *Hvem^−/−^*, *Hvem^fl/fl^* and mice with the cytoplasmic tail of *Hvem* deleted mice were bred and analyzed at La Jolla Institute for Immunology, La Jolla, CA, United States. All mice were between 7 – 24 weeks old. Animal care was in accordance with the Canadian Council on Animal Care guidelines. The studies were performed under Animal Use Protocol 215, approved by the Animal Care and Use Committee Health Sciences of the University of Alberta. Mice were housed under clean conventional housing conditions at the University of Alberta Health Sciences Laboratory Animal Services (HSLAS) facilities. Mouse studies carried out at La Jolla Institute for Immunology used animals bred and housed under specific pathogen–free conditions and were approved by the La Jolla institute for Immunology Animal Care and Use Committee under protocol AP00001007.

### Generation of HVEM mutant mice

Mouse HVEM cytoplasmic tail mutant (*Hvem^tm^*) mice were generated by the CRISPR/Cas9 system by injection of a sgRNA-Cas9 complex plus a donor specific single-stranded DNA (ssDNA) into C57BL/6 pronuclear embryos. All materials for the CRISPR/Cas9 system were purchased from Integrated DNA Technologies (IDT, Newark, NJ). The specific sgRNA targeted the front of the *Tnfrsf14* exon 7: 5’-AGAACAUCAAGUCAUGGGAG-3’. The ssDNA homology directed repair (HDR) template has a stop codon in the exon 7 of the *Tnfrsf14* locus: 5’-CCATAAGCATATGCCAGTTGGAACTTCCTCCCCGACCCAGTTATACCTGGAAAGGCTCCAGCTCCTTAGTCACTTAGCCTGTAACACAAGAACATCAAGTCATGGGAGAGCTGAAGCAAGAGGGGAGGGAGACGGGCACACAGCAATGAAAAACCCACATTCTGGGATTCCAGCTGTGTGATCTACCTCCCAAGTCTGAC-3’. The HDR repair did not occur, but we obtained an F0 founder that has an allele with a deletion of the whole exon 7 and a read-through out-of-frame amino acid sequence by splicing exon 6 to exon 8. The F0 founder was backcrossed to the WT C57BL/6 strain for at least 2 generations. We obtained homozygous offspring (*Hvem^WT^* and *Hvem^tm^*) by intercrossing the N2 generation of *Hvem^+/tm^* mice. The mice were generated at the RIKEN Center for Integrative Medical Sciences (IMS), Yokohama, Japan. All procedures related to strain construction were approved by the RIKEN IMS Animal Care and Use Committee. We confirmed that the tail-mutated HVEM protein was incapable of signaling in vitro and in vivo, as described previously [66]. For in vitro testing, the pGL4.32[luc2P/NF-κB-RE/Hygro] (NF-κB–driven firefly luciferase; Promega) and pRL-TK (*Renilla* luciferase as an internal control; Promega) plasmids were co-transfected with mouse *Hvem^WT^*, or the *Hvem^tm^* sequence with deletion of exon 7, or control vector (EGFP) plasmids into 293T cells by TransIT-LT1 (Mirus). One day later, transfected cells were co-cultured with transfected 293T cells expressing mouse LIGHT, CD160, or BTLA. On the next day, firefly and *Renilla* luciferase activity were measured through the Dual-Glo Luciferase Assay System (Promega) and detected by the EnVision 2104 Multimode Plate Reader (PerkinElmer). For in vivo testing, we used a hepatic injury model. Co-housed female littermates were injected with 2 µg αGalCer (KRN7000; Kyowa Kirin Research) in a total volume of 200 µl PBS by retro-orbital injection. Serum ALT activity was measured using a colorimetric/fluorometric assay kit (ab105134; Abcam) at 24 h after injection as described previously [66].

### Cell transfer, tamoxifen-induced conditional deletion, and disease criteria

Thymocytes or splenocytes containing the indicated number of SP T cells were injected into adult *Rag^−/−^* mice. Briefly, thymuses and spleens were removed from the donors and mashed with glass slides on ice to make a single-cell suspension, followed by filtration with a 70 µm cell strainer (Fisherbrand™). CD4 and CD8 SP T cells were isolated by negative selection from a single cell thymocyte suspension using EasySep™ Mouse Naïve CD4^+^ T Cell Isolation Kit (# 19765) and EasySep™ Mouse Naïve CD8^+^ T Cell Isolation Kit (# 19858), respectively, from STEMCELL Technologies (Vancouver, Canada) according to manufacturer’s instructions. The purity of the sorted cell population was > 96%. Viability was assessed by trypan blue exclusion and > 90% viable cells were used for experiments. Where indicated, TCR^+^ CD24^low^ cells were sorted from splenocytes of six-week-old *Btla^−/−^* mice on a FACS BD influx^TM^ cell sorter (BD Biosciences). The purity of the sorted cell populations was 92%. Fetal liver cells (FLC) were used as a source of hematopoietic stem cells (HSC) and were harvested from embryonic day 14 – 16 fetuses. A single-cell suspension was made on ice by gently pipetting the fetal livers and filtration through a 70 µm cell strainer (Fisherbrand™). Viability was assessed by trypan blue exclusion and > 90% viable cells were used for experiments. Six-eight-week-old male and female *Rag^−/−^* mice were used as recipients, and each recipient received 10 - 20 x 10^6^ FLC, followed by tamoxifen injection as described below. Some HSC recipients also received agonist anti-BTLA antibody (6A6; 10μg/g body weight) or hamster IgG isotype control antibody once per week beginning 18 days post fetal liver injection. To induce BTLA or PD-1 deletion, B6*^Cre/ERT2+/−^ Btla^fl/fl^* mice or adult *Rag^−/−^* recipients of B6*^Cre/ERT2+/−^ Btla^fl/fl^* or B6*^Cre/ERT2+/−^ Pdcd1^fl/fl^* FLC or thymocytes were intraperitoneally injected with 1.4 mg tamoxifen (Sigma-Aldrich) in corn oil [+5% (vol/vol) ethanol] on days 0, 1, 3, 5, and 6. Recipients of control B6*^CreERT2+/−^* FLC or thymocytes also received tamoxifen injection. Signs of disease included loss of weight, hunched appearance, piloerection, diarrhea, and dermatitis. Recipient mice were no longer considered disease-free when two or more of the above signs were evident, or mice had lost ≥ 20% of baseline body weight. In addition, for mice to be classified as diseased, disease signs must persist for at least two weeks. Immunohistochemistry staining was performed on tissues from multiple organs collected from recipient mice.

### Immunohistochemistry

Mice were euthanized and transcardially perfused with phosphate buffer saline (PBS), followed by 4% paraformaldehyde (PFA) in PBS. Harvested non-lymphoid organs (heart, kidneys, liver, and lungs) were immersed in 4% PFA in PBS overnight at 4°C and then transferred into fresh 30% sucrose in PBS every day for two consecutive days at 4°C. Tissues were embedded in optimum cutting temperature compound (TissueTek OCT, Sakura Finetek, 4583), frozen on liquid nitrogen, and cryosectioned (Leica CM1950) at –20°C with a thickness of 5 or 10 µM on glass slides. Following three washes in 1x PBS, tissue sections were blocked in 10% normal goat serum for 1 h at room temperature and incubated in rat anti-mouse CD4 (1:200; MCA2691; Bio-Rad) or CD8a (Biolegend, San Diego, CA) antibody overnight at 4°C. Slides were washed in PBS-Tween (0.5% Tween 20 in 1x PBS) and incubated in goat anti-rat IgG Alexa Fluor® 488 (1:200; A11006; Life Technologies) antibody for 45 min at room temperature. To visualize cellular nuclei, tissue sections were counterstained with VECTASHIELD mounting medium with DAPI (Vector Laboratories, H-1200) (42,43). Spleens from *Rag^−/−^* and WT mice were used as the CD4 negative and positive control, respectively. The negative control for primary antibody specificity was omitting the primary antibody in the staining. Immunofluorescence images were taken using either a Leica DM IRB Microscope and Open Lab software or an Axioplan, Axiovision 4.1 software (Carl Zeiss, Toronto, ON). An average of three images per section were examined.

### Flow cytometry and BrdU incorporation

Fluorochrome conjugated antibodies for flow cytometry staining used in this study were purchased from ThermoFisher Scientific: murine anti-TCRβ (H57-597), CD4 (RM4-5), CD5 (53-7.3), CD8α (53-6.7), CD44 (IM7), CD62L (MEL-14), FoxP3 (FJK-16s), BTLA (6F7), PD-1 (J43); or BioLegend: CD19 (6D5), HVEM (HMHV-1B18). GFP expression was also analyzed in mice expressing the GFP transgene. Peripheral blood samples, thymocytes, and splenocytes were stained after incubation with FcR block, which was a cocktail of anti-CD16/32 antibody (2.4G2; Bio Express, West Lebanon, NH) and mouse, rat, and hamster sera. Staining was done at 4°C for 20 minutes, followed by washing and resuspension in Hanks’ Balanced Salt Solution (HBSS) supplemented with 2% fetal bovine serum (FBS). A BD LSR II (BD Biosciences) with FlowJo software was used for data acquisition and analyses. To assess proliferation in vivo, experimental mice were treated with 2 mg BrdU in PBS by i.p. injection. BrdU incorporation was assessed in splenic T cells 24 hours after injection using a BrdU flow cytometry kit (BD PharMingen^TM^).

### Statistical analysis

Statistical analysis was performed using GraphPad Prism software. Data of biological replicates are depicted as mean ± standard error. Data were statistically analyzed using Student’s t-test with Welch’s correction or Wilcoxon matched-pairs signed rank test or Mann-Whitney test. Statistical analysis in experiments with one variable and three groups was performed using one-way ANOVA with Dunn’s multiple comparisons test or Kruskal-Wallis test (\**p* < 0.05; \*\**p* <0.01; \*\*\**p* <0.001; \*\*\*\**p* <0.0001). Disease onset/incidence was compared by the Kaplan–Meier method. Probability values reported for survival curve comparisons were calculated using the Mantel-Cox method.

## Acknowledgments

We thank Perveen Anwar, Joaquín López-Orozco and Sudip Subedi for technical assistance, and Jiaxin Lin, Kevin Zhan, Nevil Singh, and John Šedý for helpful discussions.

## Author Contributions

**Conceptualization:** Colin C. Anderson, Adeolu O. Adegoke, Troy A. Baldwin, Mitchell Kronenberg

**Formal Analysis:** Adeolu O. Adegoke, Govindarajan Thangavelu, Ting-Fang Chou, Marcos Petersen, Kevin Joannou, Qingyang Wang

**Funding Acquisition:** Colin C. Anderson, Troy A. Baldwin, Peter A. Bretscher, Hilde Cheroutre, Mitchell Kronenberg

**Investigation:** Adeolu O. Adegoke, Govindarajan Thangavelu, Ting-Fang Chou, Marcos Petersen, Julia F. May, Kevin Joannou, Qingyang Wang, Kristofor K. Ellestad

**Methodology:** Kiyokazu Kakugawa, Hilde Cheroutre

**Resources:** Kiyokazu Kakugawa, Hilde Cheroutre, Louis Boon

**Project Administration:** Colin C. Anderson

**Supervision:** Colin C. Anderson, Troy A. Baldwin, Mitchell Kronenberg, Peter A. Bretscher

**Writing – original draft:** Colin C. Anderson, Adeolu O. Adegoke, Govindarajan Thangavelu

**Writing – review & editing:** All authors

## Abbreviations

DN: double negative
DP: double positive
SP: single positive
WT: wild type
RTE: recent thymic emigrants
FLC: fetal liver cells

