## supplemental information for "Internal regulation between constitutively expressed T cell coinhibitory receptors BTLA and CD5 and tolerance in recent thymic emigrants"

**Supplementary Methods**

CD4 SP T cells were isolated by negative selection using the EasySep™ Mouse Naïve CD4+ T Cell Isolation Kit (# 19765, STEMCELL Technologies, Vancouver, Canada) according to the manufacturer's instructions, from single cell thymocyte suspensions prepared from B6.*Btla* WT and B6.*Btla*$^{-/-}$ mice (aged 8-12 weeks). TCR sequencing was conducted using the 10x Genomics V(D)J workflow. Briefly, single-cell TCR libraries were prepared using the Chromium Single Cell Mouse TCR Amplification Kit (Catalog #1000254). Libraries were pooled to achieve desired quantities for appropriate sequencing depths, as recommended by 10x Genomics, and were sequenced on an Illumina NovaSeq 6000 (v1.5) instrument. Alignment of reads was performed using the prebuilt Cell Ranger v7.2.0 mouse reference GRCm38 v7.0.0. Reads mapping and contig annotations were conducted using the 10x Genomics Cell Ranger 7.2.0 vdj pipeline via 10x Genomics Cloud Analysis (https://www.10xgenomics.com/, accessed 2024-April-10). Across samples of WT and *Btla*$^{-/-}$, a total of 4040 and 5077 T cells were sequenced, respectively.

Single-cell TCR repertoires were analyzed using scRepertoire package v2.0.0 [1] within the R environment (v4.3.1). The analysis included the examination of V gene usage, CDR3 amino acid (AA) length, and diversity along the residues of the CDR3 AA sequence, based on a Shannon score. For V gene usage we plotted the subgroup of TCRAV and TCRBV genes [2,3] for a better visualization. Shared and unique clonotypes, defined by their V(D)J gene usage, in WT and *Btla*$^{-/-}$ mice were depicted by Venn diagrams. The top 20 clones from each sample were visualized using alluvial plots. Clones were represented as stacked bins, with their height indicating their frequency in the sample. Shared clones between samples are linked.

Acknowledgement: Libraries were created at the Advanced Cell Exploration Core at the University of Alberta Faculty of Medicine & Dentistry, RRID:SCR_019182, which receives financial support from the Faculty of Medicine & Dentistry, the Li Ka Shing Institute of Virology, Striving for Pandemic Preparedness – The Alberta Research Consortium, and Canada Foundation for Innovation (CFI) awards to contributing investigators. Sequencing was done at the University of Calgary's Centre for Health Genomics and Informatics.

1. Borcherding N, Bormann NL, Kraus G. scRepertoire: An R-based toolkit for single-cell immune receptor analysis. F1000Research. 2020;9: 47. doi:10.12688/f1000research.22139.2

2. Bosc N, Lefranc M-P. The mouse (Mus musculus) T cell receptor alpha (TRA) and delta (TRD) variable genes. Dev Comp Immunol. 2003;27: 465–497. doi:10.1016/s0145-305x(03)00027-2

3. Bosc N, Lefranc M-P. The Mouse (Mus musculus) T Cell Receptor Beta Variable (TRBV), Diversity (TRBD) and Joining (TRBJ) Genes. Exp Clin Immunogenetics. 2000;17: 216–228. doi:10.1159/000019141

### Supplementary Figures

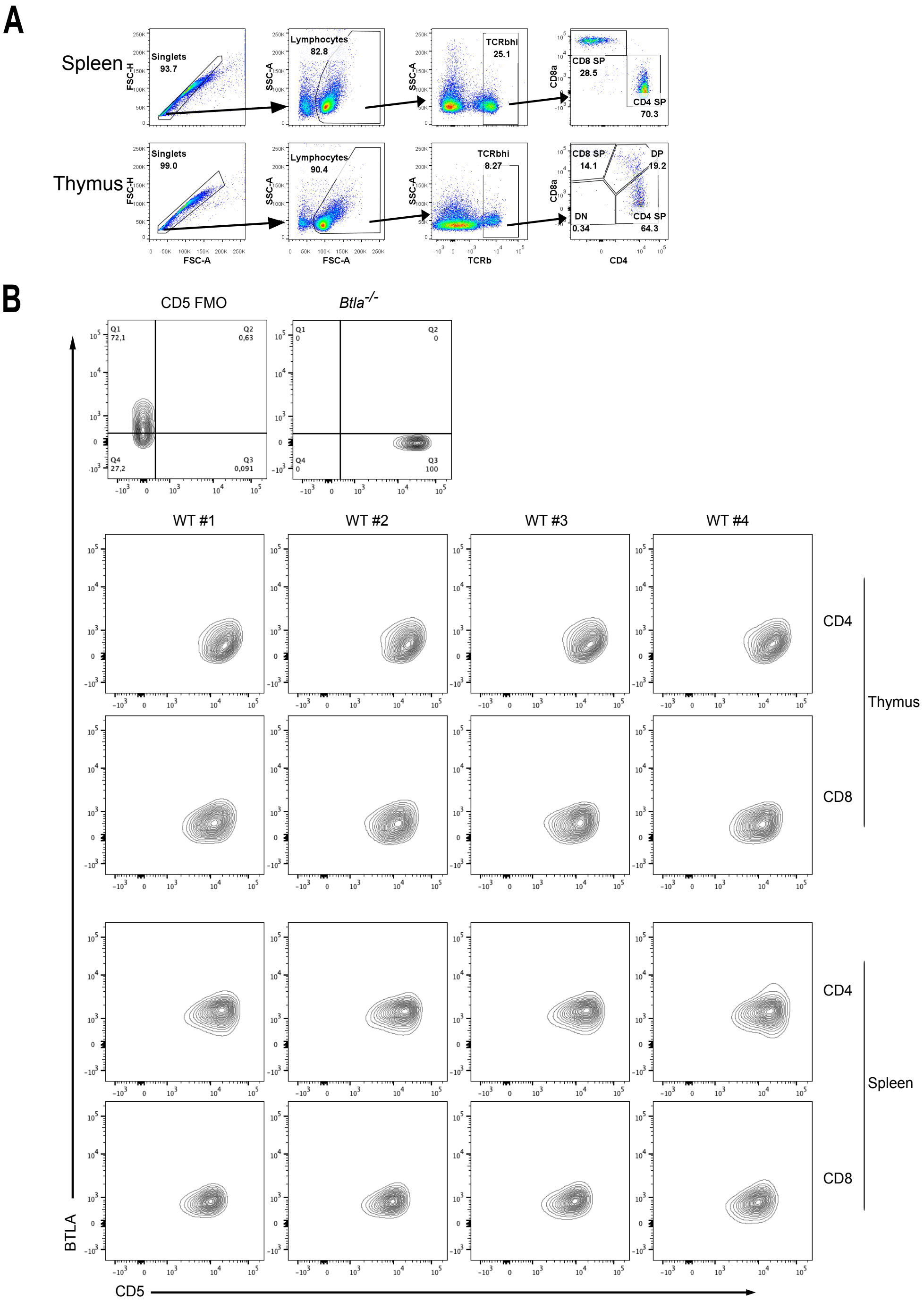


**S1 Fig. Gating used for analysis of coinhibitory expression and lack of inverse relationship between CD5 and BTLA expression within T cell subsets of each tissue.** [**A**] Gating strategy for CD5 and BTLA in the spleen and thymus. We gated on single cells, followed by lymphocytes, and then the TCRβ$^{hi}$ SP T cells in the spleen (upper row) and thymus (lower row). [**B**] BTLA and CD5 expression in thymic or splenic CD4 and CD8 SP T cells of four WT B6 mice; top row shows the CD5 FMO staining (left) and staining of *Btla*$^{-/-}$ cells (right).

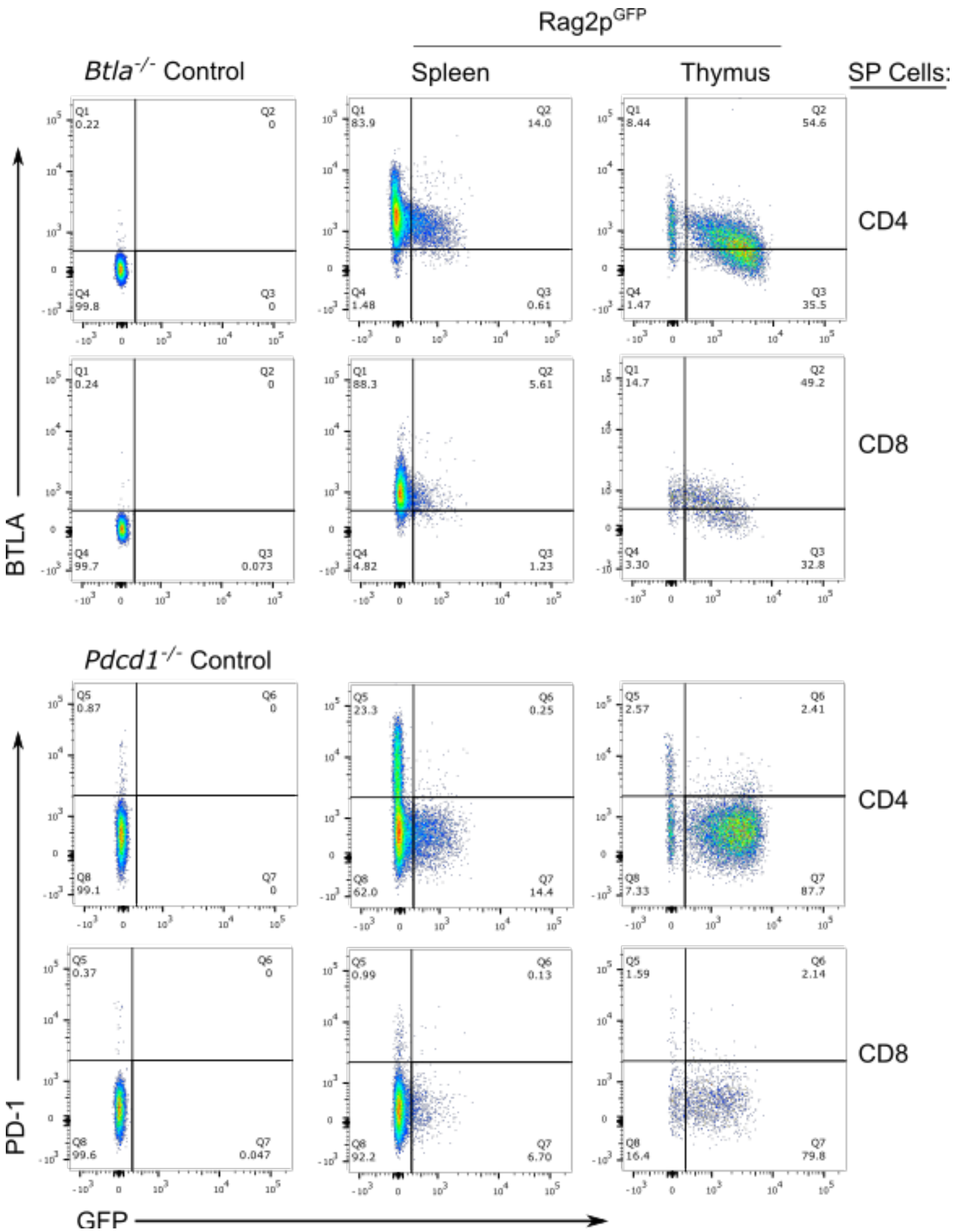


**S2 Fig.** BTLA and PD-1 expression was examined in newly generated single positive (SP) T cells ($GFP^+$) and established SP T cells ($GFP^-$) in the spleen and thymus of *Rag2p^GFP^* mice (gated on $TCR\beta^+$ and either $CD4^+$ $CD8^-$ or $CD4^-$ $CD8^+$ cells; representative of n = 4). Staining of similarly gated splenocytes from control *Btla*$^{-/-}$ and *Pdcd1*$^{-/-}$ mice is shown.

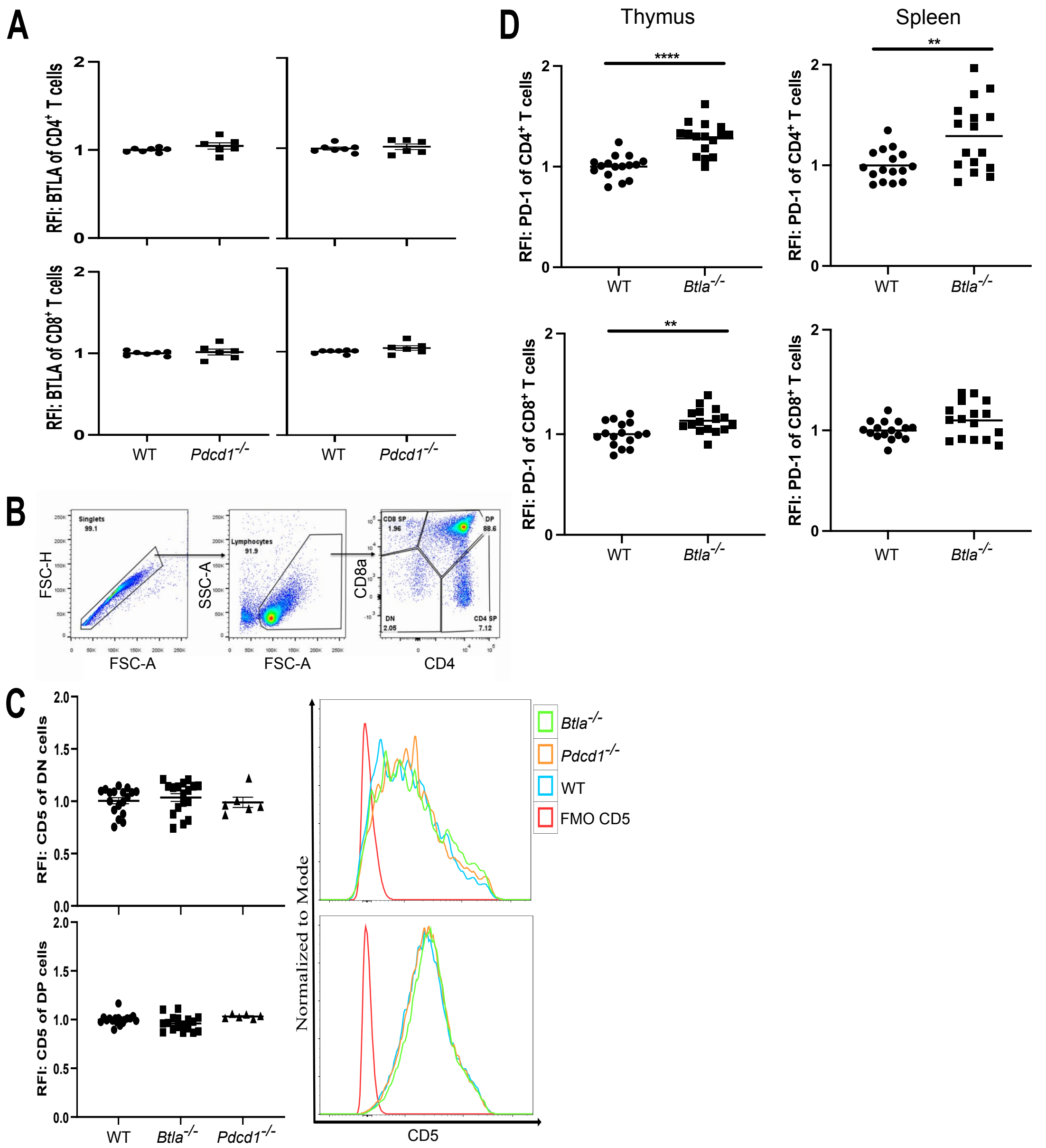


**S3 Fig. PD-1 deficiency does not affect BTLA expression and BTLA deficiency does not affect CD5 expression in DN or DP thymocytes but does increase PD-1 expression in thymic and splenic SP T cells.** [**A**] RFI of BTLA in thymic (left column) or splenic T cells (right column) of WT (n = 7; *B6.Foxp3*$^{GFP}$) and PD-1$^{-/-}$ (n = 6; *B6.Foxp3*$^{EGFP}$ x *Pdcd1*$^{-/-}$) mice. To calculate RFI, data were normalized to the average Btla MFI of the WT splenic or thymic SP T cells in each individual experiment. Dots indicate individual mice from two experiments. [**B, C**] CD5 expression in developing DN and DP T cells is not increased in the Btla$^{-/-}$ or PD-1$^{-/-}$ mice. [**B**] Gating strategy for CD5 expression in the immature T cells showing gating on single cells, followed by lymphocytes, and then the DP and DN cells in the thymus. [**C**] RFI of CD5 in DN and DP cells in the WT, *Btla*$^{-/-}$, *Pdcd1*$^{-/-}$ mice (left) and representative histograms (right). [**D**] RFI of PD-1 in thymic (left column) or splenic T cells (right column) of WT (n = 16; B6.*Foxp3*$^{GFP}$ and B6.*Nur77*$^{GFP}$) and *Btla*$^{-/-}$ (n = 16; B6.*Foxp3*$^{GFP}$*Btla*$^{-/-}$ and B6.*Nur77*$^{GFP}$*Btla*$^{-/-}$) mice. To calculate RFI, data were normalized to the average MFI of CD5 or PD-1 of the WT thymic or splenic T cells in each individual experiment. Dots indicate individual mice from a minimum of three independent experiments. *$p < 0.05$, **$p < 0.01$, ****$p < 0.0001$.

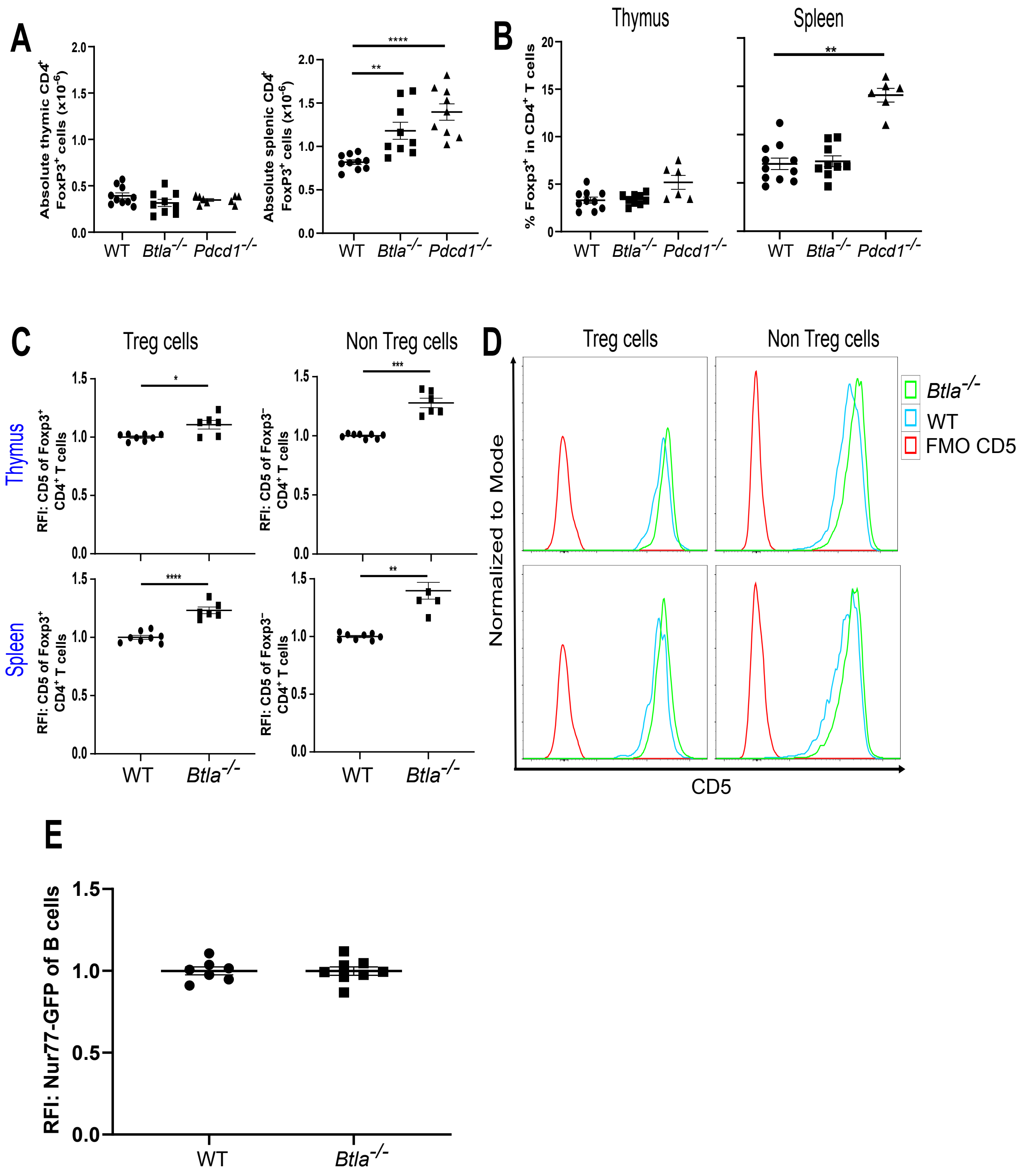


**S4 Fig.** Comparison of the absolute count [**A**] and percent [**B**] of CD4$^+$ Foxp3$^+$ T cells in the thymus and spleen of adult B6 WT, *Btla*$^{-/-}$, and *Pdcd1*$^{-/-}$ mice. [**C**] RFIs of CD5 in Treg and non-Treg cells of WT and *Btla*$^{-/-}$ mice, and representative histograms [**D**]. To calculate RFIs, data are normalized to the average MFI of CD5 of the WT thymic or splenic T cells in each individual experiment. [**E**] RFI of Nur77$^{GFP}$ in splenic B cells of WT (n = 7) and *Btla*$^{-/-}$ (n = 8) mice. To calculate RFI, data were normalized to the average Nur77-GFP MFI of the WT splenic B cells in each individual experiment. Dots indicate individual mice from two [E] or a minimum of three [A – D] independent experiments. *$p < 0.05$, **$p < 0.01$, ****$p < 0.0001$.

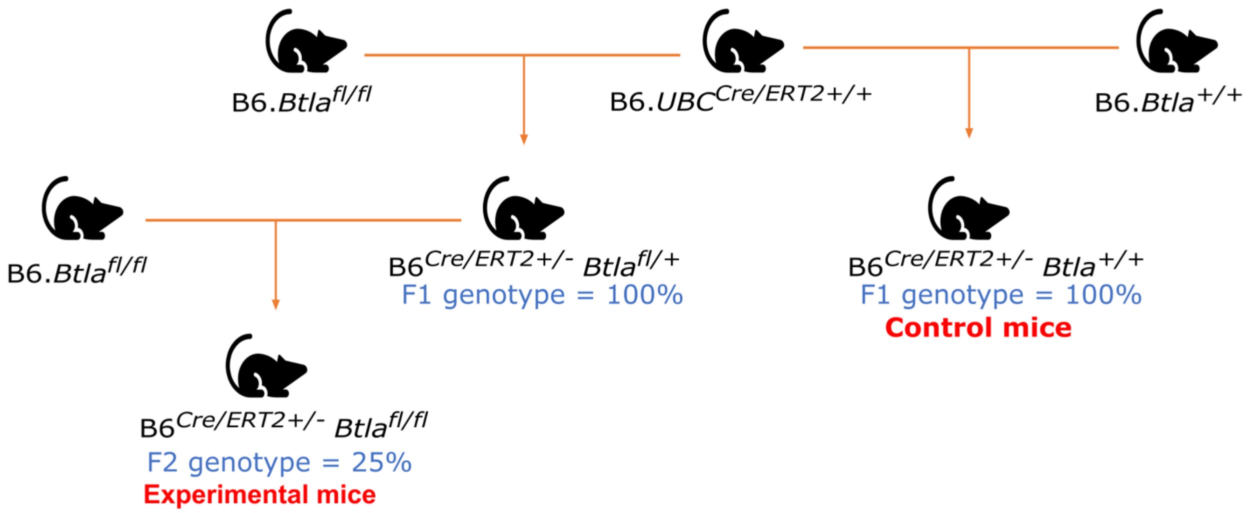


**S5 Fig.** *Breeding strategy for the B6.UBC$^{Cre/ERT2+}$Btla$^{fl/fl}$ mice.* The B6.*Btla*$^{fl/fl}$ strain was crossed to a tamoxifen-inducible Cre recombinase expressing strain, B6.*UBC*$^{Cre/ERT2+/+}$ (abbreviated B6$^{Cre/ERT2+/+}$). The resulting F1 B6.*UBC*$^{Cre/ERT2+/-}$*Btla*$^{fl/-}$ was backcrossed to B6.*Btla*$^{fl/fl}$ to generate B6.*UBC*$^{Cre/ERT2+/-}$*Btla*$^{fl/fl}$. For the control WT mice, B6.*UBC*$^{Cre/ERT2+/+}$ mouse was crossed to B6.*Btla*$^{+/+}$ mice to generate B6.*UBC*$^{Cre/ERT2+/-}$*Btla*$^{+/+}$ mice.

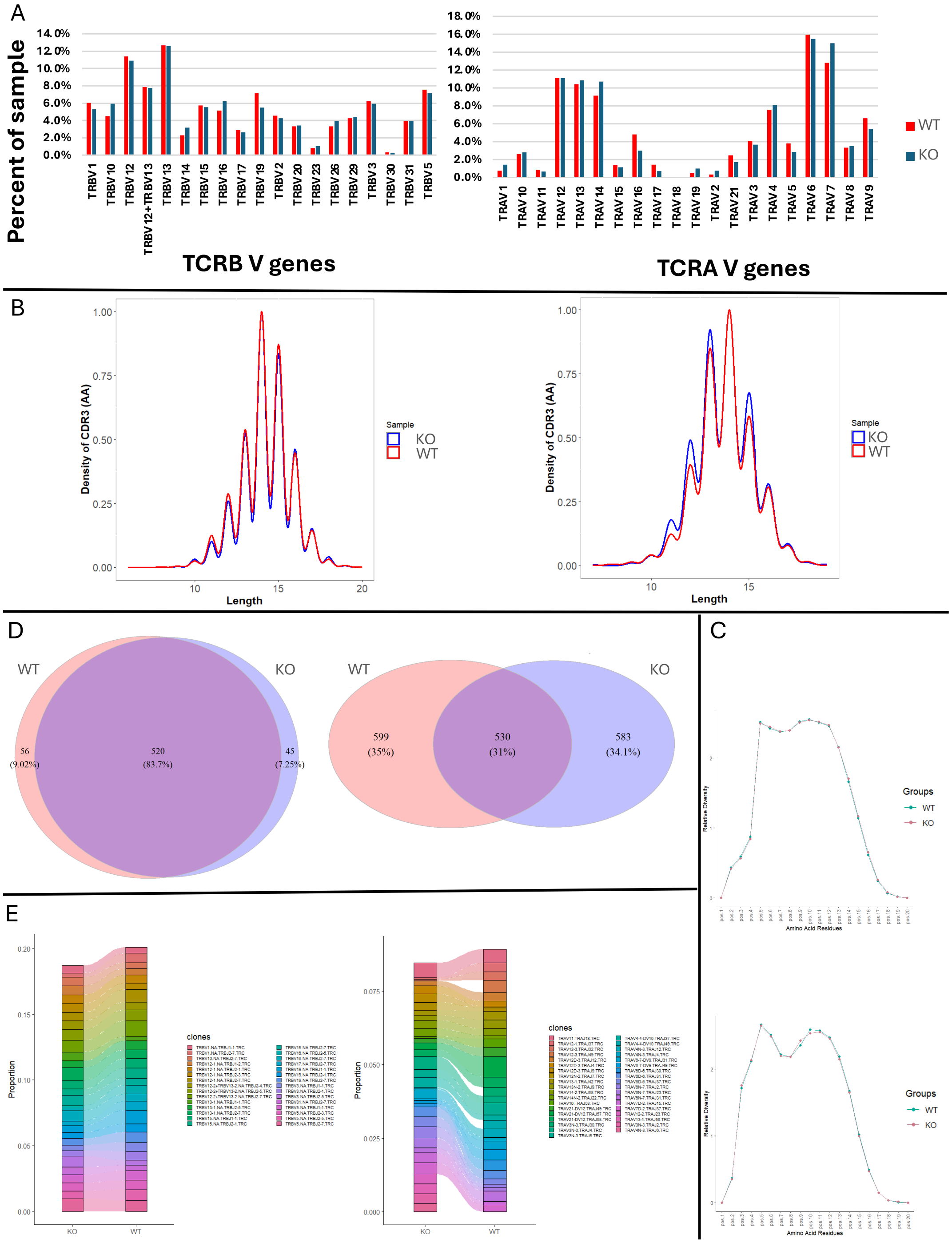


**S6 Fig. TCR Repertoire Analysis of β- and α-Chain Using Single-Cell RNA Sequencing of Sorted Thymic CD4 SP T Cells from B6 WT and B6.*Btla*$^{-/-}$ (KO) Mice.** [A] Transcript based assessment of relative TCRβ (left) and TCRα (right) V family gene usage by sample. [B] Length distribution of the CDR3 amino acid (AA) sequences for β-chain (left) and α-chain (right). [C] Diversity along the residues of the CDR3 AA sequence for the β-chain (top) and α-chain (bottom). [D] Venn diagrams depicting shared and non-shared TCRβ clonotypes (left) and TCRα clonotypes (right), defined by their V(D)J gene usage. [E] Alluvial plots illustrating the 20 most frequent clones for the β-chain (left) and α-chain (right). Shared clones are linked between samples.

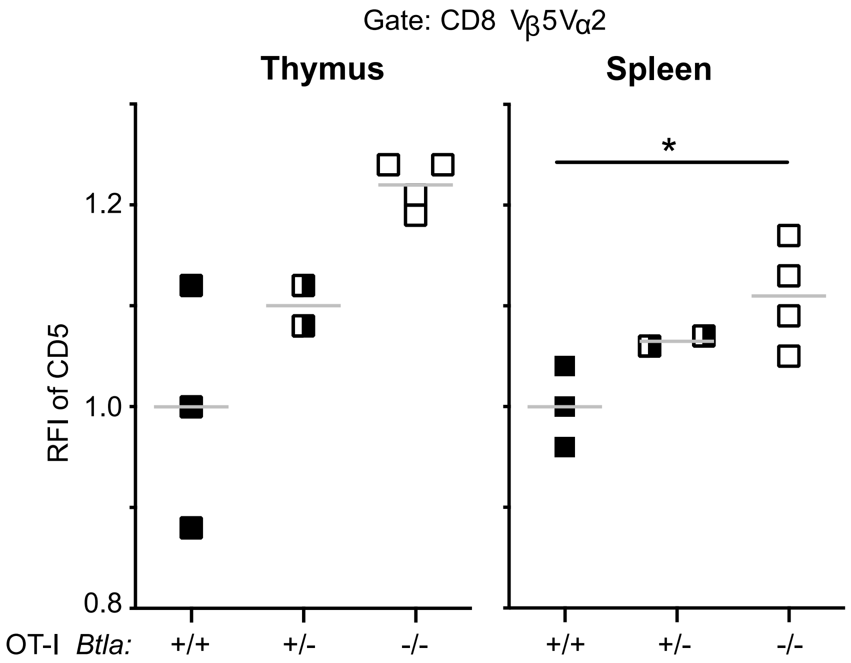


**S7 Fig. BTLA deficiency increases CD5 levels in TCR transgenic OT-I CD8 T cells.** RFI of CD5 in all $CD8^{+}$ TCR-$V\beta5V\alpha2^{+}$ cells in the thymus (left) and spleen (right) of OT-I.$Btla^{+/+}$ (WT OT-I; n = 3), OT-I.$Btla^{+/-}$ (n = 2) and OT-I.$Btla^{-/-}$ mice (n = 4). The grey line is the mean of the RFI. CD5 expression on splenic OT-I.$Btla^{+/+}$ T cells was significantly lower than on BTLA deficient ($Btla^{-/-}$) OT-I T cells, $*p < 0.05$.

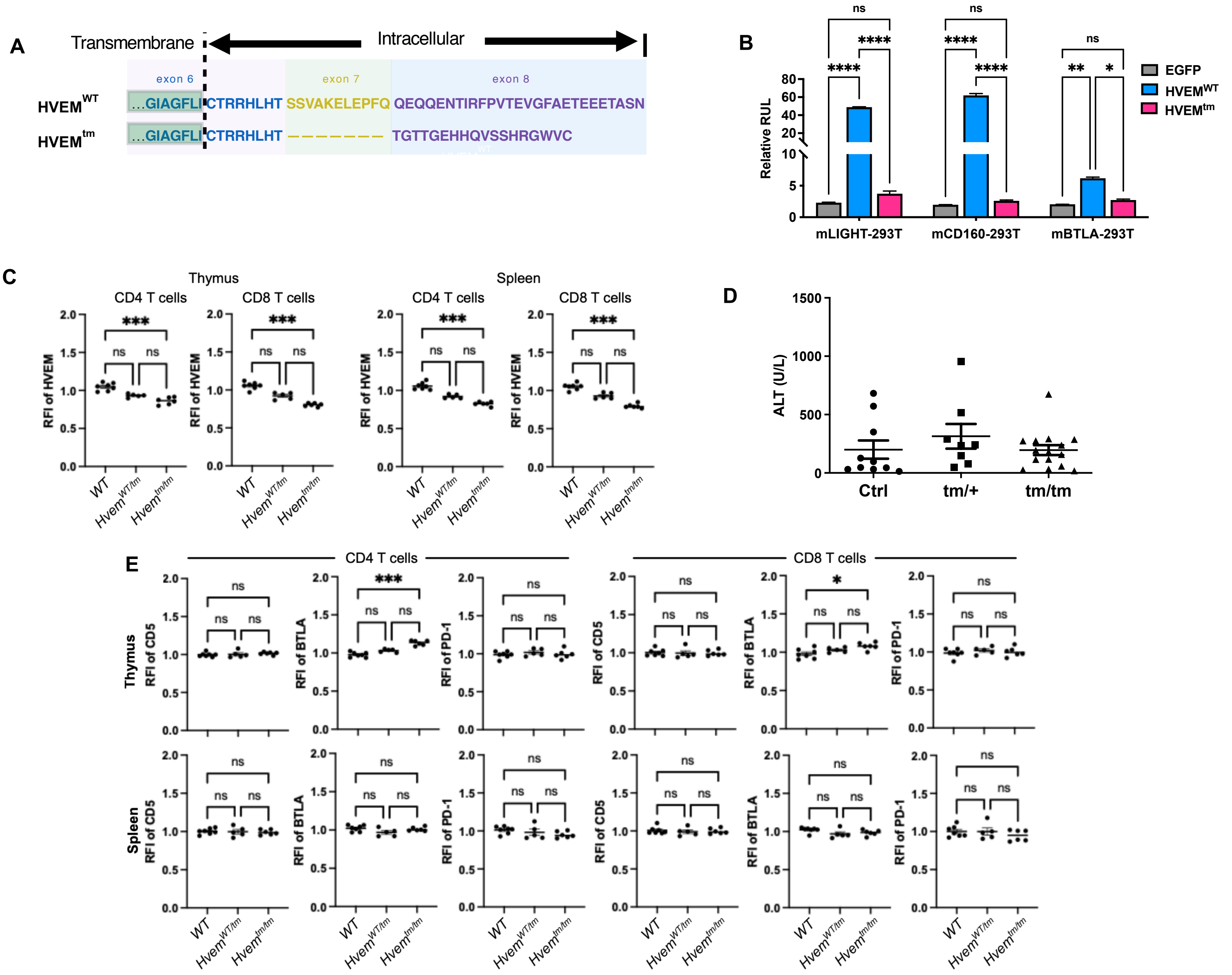


**S8 Fig. HVEM dependent control of CD5 and PD-1 expression does not require HVEM signaling.** [**A**] Schematic of amino acid sequences of the HVEM intracytoplasmic regions in WT and HVEM cytoplasmic tail mutant (HVEM$^{tm}$) mice. [**B**] The HVEM$^{tm}$ construct does not signal. 293T cells were co-transfected with mouse HVEM$^{WT}$, HVEM$^{tm}$, or control EGFP vector along with a NF-κB–driven luciferase (NF-κB-Luc) vector. mHVEM/NF-κB-Luc–expressed cells were co-cultured with LIGHT, CD160, or BTLA-transfected 293T cells as indicated. Luciferase activity was measured after 18 h. RLU (relative light units) is the ratio of Firefly luciferase luminescence to Renilla luciferase luminescence. Data shown are mean ± SEM (n = 2) and depict representative results from two independent experiments. *$p < 0.05$; **$p < 0.01$; **** $p < 0.0001$ for two-way ANOVA. [**C**] Thymocytes and splenocytes from the indicated strains, gated on SP CD4 and SP CD8 T cells, were examined by flow cytometry for expression of HVEM (n = 5 – 7 mice per group). ***$p < 0.001$ [**D**] Co-housed littermates were injected with 2 µg αGalCer by the retro-orbital route. Serum ALT activity was measured at 24 h from the indicated mice. Data shown are mean ± SEM. One-way ANOVA (n = 8 – 15 mice per group). [**E**] Thymocytes and splenocytes from the indicated strains, gated on SP CD4 and SP CD8 T cells, were examined by flow cytometry for expression of CD5, BTLA, and PD-1. Data represent pooled results from at least two independent experiments (n = 5 – 7 mice per group). *$p <$ 0.05, ***$p <$0.001.

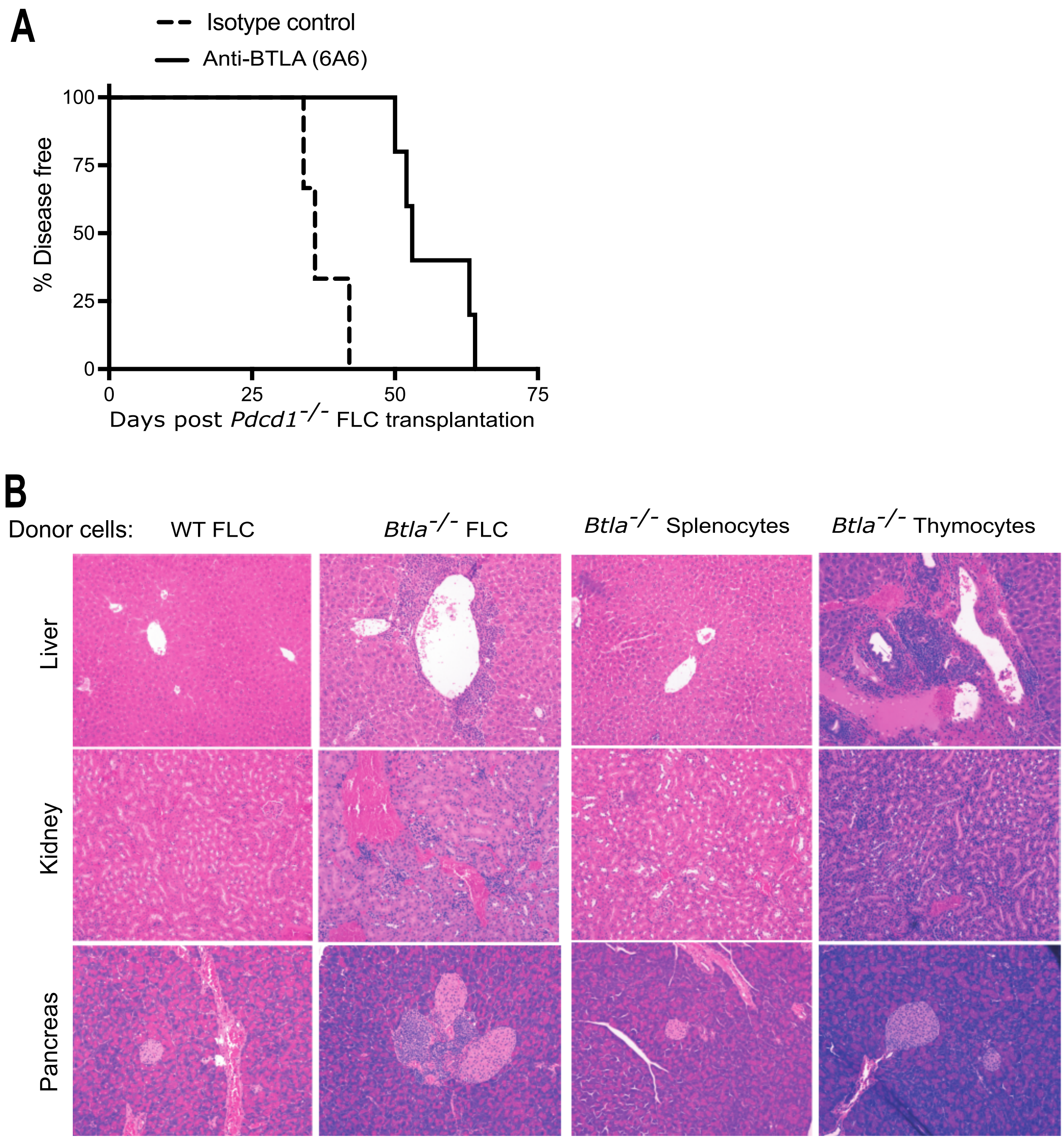


**S9 Fig.** [**A**] Adult *Rag*$^{-/-}$ recipients were given *Pdcd1*$^{-/-}$ FLC followed by treatment with anti-BTLA (n = 5) or isotype control antibody (n=3); $p = 0.004$. [**B**] *Rag*$^{-/-}$ mice (shown in Fig. 6A) were given WT FLC, *Btla*$^{-/-}$ FLC, *Btla*$^{-/-}$ splenocytes or *Btla*$^{-/-}$ thymocytes. Representative histology (hematoxylin and eosin staining) of the liver, kidney and pancreas of the recipients (n = 4-6/group) 60-65 days post cell transfer. The frequency of substantial infiltration of liver, kidney, or pancreas was 4, 3, and 2 out of 5 mice, respectively, for recipients of *Btla*$^{-/-}$ FLC. Infiltration was not seen in the organs of recipients of WT FLC or *Btla*$^{-/-}$ splenocytes.

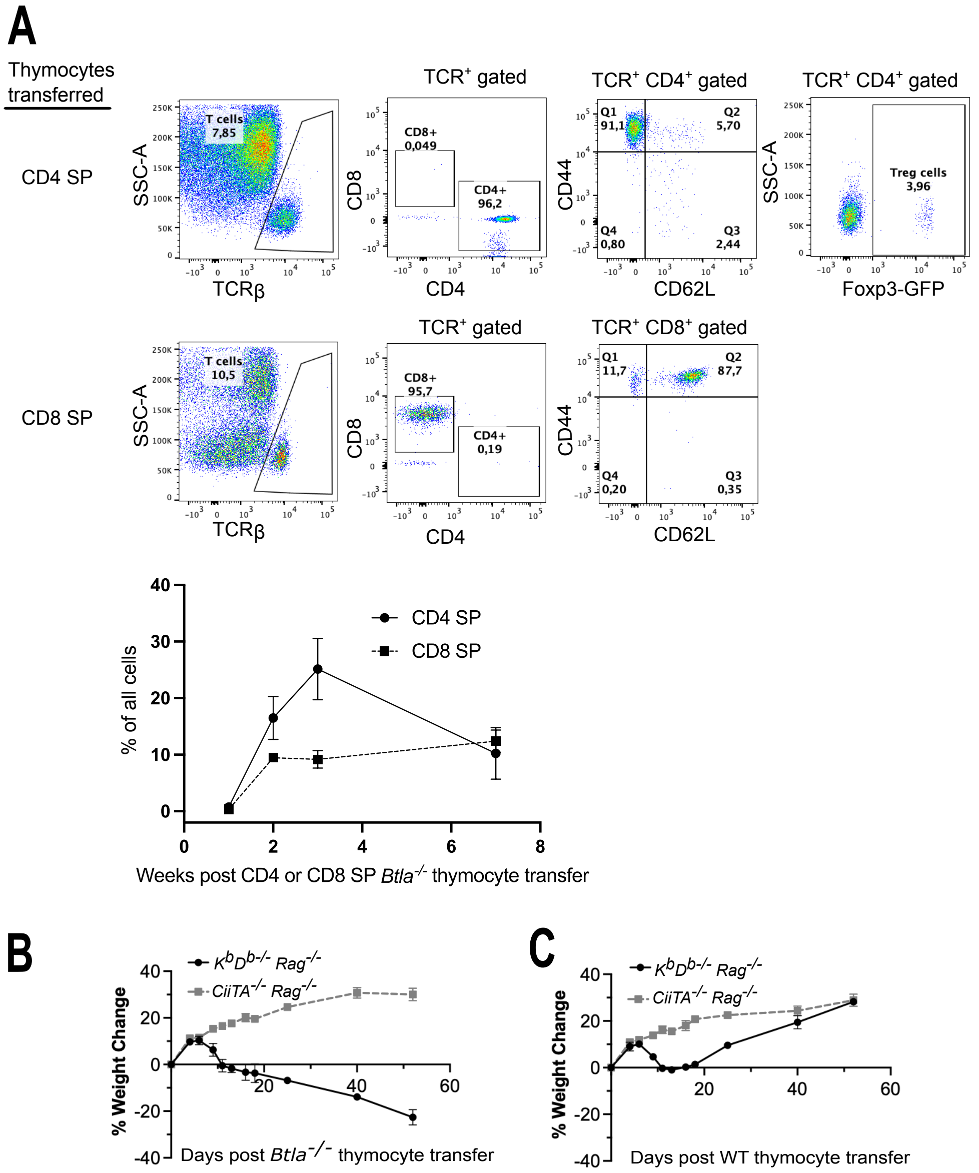


**S10 Fig. T cells post transfer of sorted B6.Foxp3$^{EGFP}$ *Btla*$^{-/-}$ thymocytes to *Rag*$^{-/-}$ recipients.** [**A**] Top: Representative flow cytometry analysis of peripheral blood at 7 weeks post transfer of sorted CD4 or CD8 SP *Btla*$^{-/-}$ thymocytes given to *Rag*$^{-/-}$ mice. Bottom: Frequency (mean ± SE, n = 4 = 5) of the indicated T cell subset within peripheral blood cells over time; disease analysis of these mice is shown in Fig. 7A. [**B,C**] *Btla*$^{-/-}$ *CD4 T cells cause disease in MHC class I deficient recipients while Btla*$^{-/-}$ *CD8 T cells do not cause disease in MHC II deficient recipients.* We adoptively transferred 3 x $10^6$ MACS-sorted CD4 SP or CD8 SP thymocytes i.v. to 8 – 10-wk old $K^bD^{b-/-}$ *Rag*$^{-/-}$ mice or *CiiTA*$^{-/-}$ *Rag*$^{-/-}$, respectively. [**B**] Weight change post cell transfer in recipients (n = 4/group) of B6.*Foxp3*$^{EGFP}$ *Btla*$^{-/-}$ cells. [**C**] Weight change post cell transfer in recipients (n = 3 - 4/group) of B6.*Foxp3*$^{EGFP}$ (WT) cells. Mice monitored for several weeks or until after losing ≥ 20% of baseline body weight, whichever came first.

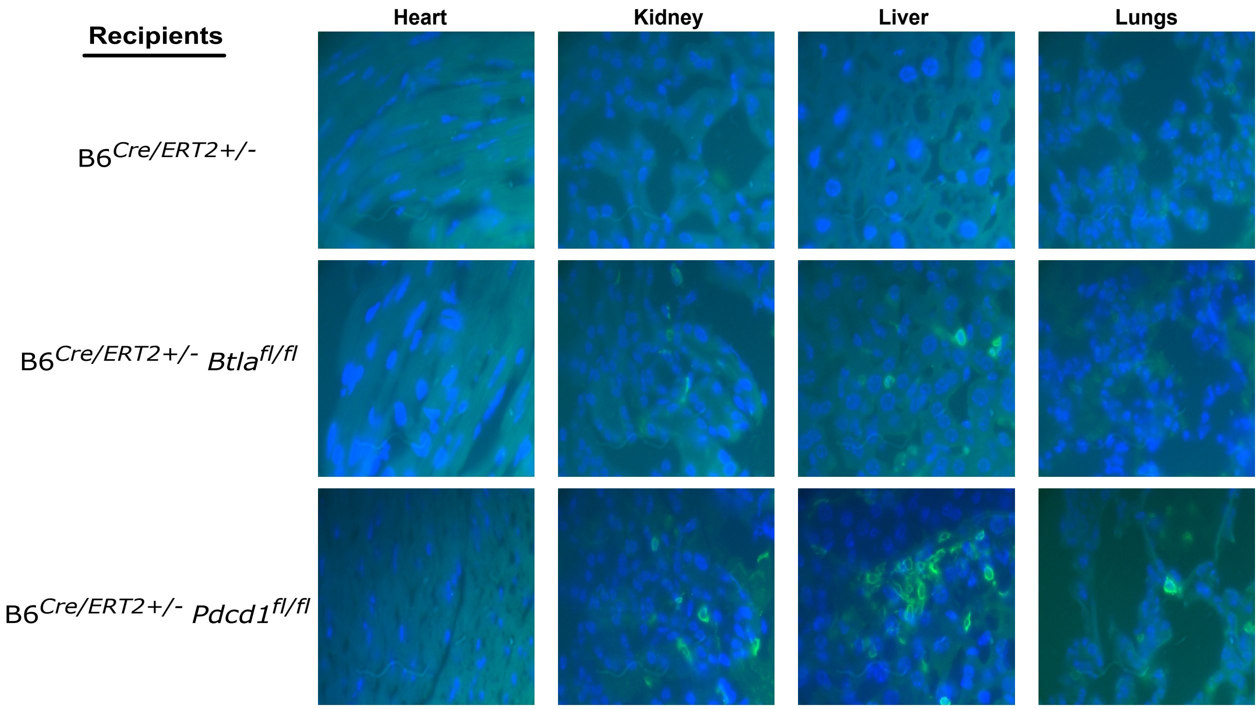


**S11 Fig. Representative immunofluorescence staining (original magnification x 1000) of heart, kidney, liver, and lungs of individual recipients of B6$^{Cre/ERT2+/–}$, B6$^{Cre/ERT2+/–}$ *Btla*$^{fl/fl}$, and B6$^{Cre/ERT2+/–}$ *Pdcd1*$^{fl/fl}$ thymocytes.** Blue: staining with the nuclear marker 4',6'-diamidino-2-phenylindole (DAPI); green: CD4 staining. CD4 SP T cell infiltration was found in the kidneys and livers of B6$^{Cre/ERT2+/–}$ *Btla*$^{fl/fl}$ thymocyte recipients (n = 9) and the kidneys, livers, and lungs of B6$^{Cre/ERT2+/–}$ *Pdcd1*$^{fl/fl}$ thymocyte recipients (n = 4). In contrast, no infiltration of CD4 SP T cells was seen in the hearts of B6$^{Cre/ERT2+/–}$ *Btla*$^{fl/fl}$ or B6$^{Cre/ERT2+/–}$ *PD-1*$^{fl/fl}$ thymocyte recipients. There was no CD4 SP T cell infiltration in the examined organs of the control B6$^{Cre/ERT2+/–}$ thymocyte recipients (n = 9).
